# Deciphering Cell Fate and Clonal Dynamics via Integrative Single-Cell Lineage Modeling

**DOI:** 10.1101/2025.09.01.673503

**Authors:** Yuntian Fu, Divij Mathew, Mingshuang Wang, Xinyi E. Chen, Kevin Z. Lin, Dylan Schaff, Sydney M. Shaffer, Drew M. Pardoll, Christina Jackson, Nancy R. Zhang

**Affiliations:** Graduate Program in Genomics and Computational Biology, University of Pennsylvania Perelman School of Medicine, Philadelphia, PA, USA; Department of Systems Pharmacology and Translational Therapeutics, Perelman School of Medicine, University of Pennsylvania, Philadelphia, PA, USA; Institute for Immunology and Immune Health, Perelman School of Medicine, University of Pennsylvania, Philadelphia, PA, USA; Parker Institute for Cancer Immunotherapy, Perelman School of Medicine, University of Pennsylvania, Philadelphia, PA, USA; Department of Neurosurgery, Perelman School of Medicine, University of Pennsylvania, Philadelphia, PA, USA; The Mark Foundation Center for Immunotherapy, Immune Signaling, and Radiation, University of Pennsylvania, Philadelphia, PA, USA; Department of Biostatistics, University of Washington, Seattle, WA, USA; Merck & Co, Incorporated, Rahway, NJ, USA; Department of Pathology and Laboratory Medicine, University of Pennsylvania, Philadelphia, PA, USA; Department of Bioengineering, University of Pennsylvania, Philadelphia, PA, USA; Bloomberg-Kimmel Institute for Cancer Immunotherapy, Baltimore, MD, USA; Mark Center for Advanced Genomics and Imaging at Johns Hopkins University, Baltimore, MD, USA; Department of Statistics, University of Pennsylvania, Philadelphia, PA, USA

## Abstract

Through natural or synthetic lineage barcodes, single-cell technologies now enable the joint measurement of molecular states and clonal identities, providing an unprecedented opportunity to study cell fate and dynamics. Yet, most computational methods for inferring cell development and differentiation rely exclusively on transcriptional similarity, overlooking the lineage information encoded by lineage barcodes. This limitation is exemplified by T cells, where subtle transcriptional differences mark divergent fates with distinct biological activity. Single-cell RNA and matched TCR sequencing is now ubiquitous in the analysis of clinical samples, where the TCR sequence provides an endogenous clonal barcode and could reveal clonal T cell responses. We present Clonotrace, a computational framework that jointly models gene expression and clonotype information to infer cell state transitions and fate biases with higher fidelity. While motivated by challenges in analyzing T cell populations, especially in the tumor microenvironment and immunotherapy settings, Clonotrace is broadly applicable to any lineage-barcoded single-cell dataset. Across diverse systems including T cells, hematopoietic differentiation, and cancer therapy resistance models, Clonotrace reveals differentiation hierarchies, distinguishes unipotent from multipotent states, and identifies candidate fate-determining genes driving lineage commitment.

## Introduction

Single cell clonal studies are now feasible at scale, where both molecular and cell lineage information are collected at single cell resolution^1–6^. Prominent examples include joint single-cell RNA-seq and TCR-seq, which are increasingly applied to T cell atlasing in diverse disease contexts, as well as lineage barcoding experiments, for which robust laboratory protocols have matured. These multimodal single cell transcriptomic and matched clonomics datasets provide rich information that links molecular state to clonal identity, offering unique insights into cellular dynamics. However, most existing computational methods for inferring dynamics, such as trajectory inference^7–10^, pseudotime ordering^11, 12^, and RNA velocity^13–15^, consider only the geometry of high-dimensional molecular profiles, ignoring lineage information. While powerful in many developmental systems, these tools often fail to recover nuanced dynamics involving nonlinear or abrupt transitions, branching paths, and reversible intermediate states.

T cells provide a clear example of the complexity: their behavior is shaped by a combination of antigen response determined by their T cell receptor (TCR) sequence, gene expression, and environmental cues including metabolic signals, cytokines, and co-stimulatory or co-inhibitory inputs. In T cells, fate boundaries are subtle and transcriptional profiles extensively overlap^16^. As a result, embeddings of single cell T cell data often lack clear low-dimensional structures amenable to trajectory modeling, and RNA velocity frequently performs poorly^17^. Clinical T cell datasets pose an additional challenge: intermediate transitional states are often underrepresented, making trajectories difficult to reconstruct. Consequently, despite the wealth of emerging T cell atlases, the dynamics of T cell in disease remain difficult to resolve.

Although reconstructing T cell dynamics solely from transcriptional information is challenging, most published and ongoing studies now profile both gene expression and paired TCR sequences at the single-cell level. The TCR serves as a natural clonal barcode, providing lineage information, yet its use in this context remains limited. Methods that have leveraged TCR information mostly focus on analyzing receptor sequence similarity to infer antigen specificity or associate receptor features with transcriptionally defined states^18–21^. Co-embedding of receptor and gene expression data has shown promise in settings with well-defined antigenic stimuli^22, 23^, but these approaches are harder to apply in complex diseases such as cancer, where antigens are diverse, patient-specific and of unknown identity. As others have shown^24^ — and as our results also confirm — cells with similar TCR sequences do not always localize in transcriptomic space, and conversely, cells with distinct receptors can exhibit highly similar transcriptional profiles. A few tools do treat TCRs as clonal identifiers: STARTRAC^25^, for example, quantifies receptor sharing between cell types with an entropy-based score, and LRT^26^ assigns clonotypes to trajectories using Dirichlet–multinomial clustering. While these approaches represent important steps, they remain tethered to transcriptome-derived partitions and thus cannot fully exploit clonal information to reveal latent structure and dynamics that lie beyond predefined annotations.

The concept of integrating transcriptomic data with clonal information also extends to the field of lineage barcoding. Recent advances in experimental lineage tracing, particularly in vitro methods using synthetic DNA barcodes^3–6, 27^, have enabled more comprehensive and precise tracking of clonal fate^1, 2, 28^. Yet, existing pipelines^29, 30^ typically depend heavily on longitudinal sampling, which is often infeasible in clinical studies, and they do not integrate the molecular and clonal modalities in a way that maximizes their joint potential.

With the growing application of single-cell RNA-seq paired with TCR profiling to clinical samples, and with the maturation of experimental lineage tracing methods, there is a pressing need for computational tools that can analyze multimodal molecular and clonal data. In this study, we introduce Clonotrace, a general framework for integrating lineage and transcriptomic information to uncover differentiation trajectories and the underlying molecular programs that drive fate decisions. A central concept in Clonotrace is the clonotype profile—the characteristic pattern of how cells from a given clone are distributed within the gene expression landscape. These patterns often recur across clones and can reveal shared differentiation behaviors or fate tendencies. Rather than relying on predefined cell types or cell trajectories, Clonotrace learns structure directly from the joint distribution of gene expression and lineage. This enables the discovery of recurrent patterns of differentiation that are not visible through gene expression alone, which makes Clonotrace well-suited to complex systems like T cells, where traditional clustering and trajectory inference often fail to resolve meaningful heterogeneity.

We applied Clonotrace to both in vitro lineage barcoding experiments and clinical single-cell RNA and TCR sequencing data from cancer immunotherapy studies. Through diverse data sets, we illustrate Clonotrace’s flexibility and the immense potential of cross-modal molecular and clonal integration: In a classic hematopoiesis system^6^, Clonotrace distinguishes between multipotent and unipotent cells, identifying cells whose progeny diverges toward multiple fates defined by distinct transcriptional programs. In an in vitro cancer treatment resistance model, Clonotrace successfully recovers temporal structure that is obscured by conventional transcriptome-only analyses. In single-cell RNAseq and TCRseq of CD8 T cells from two clinical cancer studies, the clonotype profiles inferred by Clonotrace uncover transcriptional heterogeneity not captured by static clustering. These profiles reveal subtle molecular differences that underlie key fate decisions relevant to disease progression. Applied to longitudinal data^31^ from a clinical trial of cancer patients undergoing immunotherapy, Clonotrace further reveals dynamic shifts in T cell behavior over the course of treatment, highlighting the complexity of immune responses in vivo.

## Results

### Clonotype profiles as dynamic descriptors of cell state and fate

Clonotrace is built on the central concept of the clonotype profile (Figure 1A). Each clone consists of a set of cells, and each cell’s location in the transcriptomic embedding reflects its gene expression phenotype. The distribution of a clone’s cells within this embedding captures its transcriptional behavior. While different clones may occupy distinct regions of the embedding, it is also common for multiple clones to exhibit similar embedding patterns—where their cells follow a shared distribution in gene expression space. Thus, we define a clonotype profile as a distribution in the transcriptomic embedding that is shared across multiple clones. Such shared distributions reveal conserved developmental or functional programs that manifest repeatedly across distinct clonal lineages. Importantly, clonotype profiles expose dynamic features not evident from transcriptomic data alone—for example, distinguishing whether a cell is unipotent or bipotent in its lineage potential (Figure 1B).

**Figure 1:**
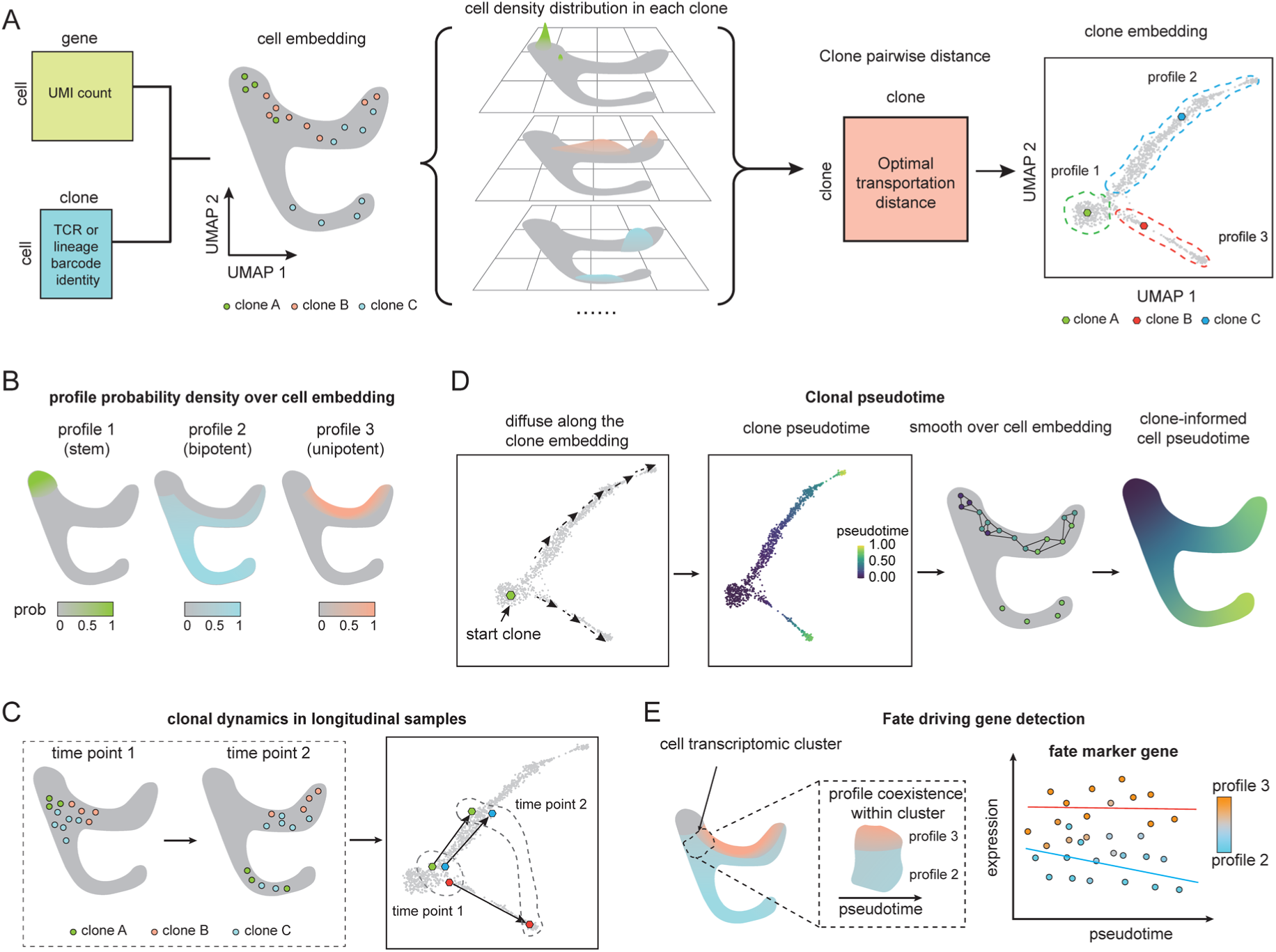
Overview of Clonotrace. **A** The scheme of clone embedding generation. **B** The cell density distribution over the cell embedding for profiles generated in the toy example in **A**. **C** An example of how clonal embedding combined with longitudinal sampling reveals longitudinal clonal dynamics. **D** The scheme for the inference of clone-informed pseudotime. **E** Comparison of profiles within the same cluster, after accounting for differences in pseudotime, yields putative fate driving genes. To differentiate between clone and cell embeddings, clone embeddings are shown with a box frame, and cell embeddings are shown without.

Clonotrace clusters clones into clonotype profiles through a two-step process. First, it constructs a “seed profile” for each clone by mapping the density of its constituent cells in transcriptomic space. Because single-cell RNA-seq data are sparse and noisy, particularly in systems like T cells where many clones are only weakly sampled due to low intrinsic cytoplasmic volume and overall mRNA content, Clonotrace smooths each clone’s footprint over a nearest-neighbor graph constructed from the full cell population (Supplementary Figure 1 and 2). This generates a continuous representation of each clone’s transcriptional distribution, mitigating sampling variability and allowing the incorporation of contextual information from non-expanded clones. Next, to quantify the similarity between clones, Clonotrace computes pairwise optimal transport distances between their smoothed density profiles. The resulting optimal transport distance matrix is embedded into a low-dimensional space, in which clones with similar distributions are placed nearby. As we will show in the ensuing examples, this clone embedding offers a clearer and more interpretable summary of clonal dynamics than the raw transcriptomic embedding, which can obscure such patterns due to sparse sampling. In longitudinal datasets, where samples are collected across multiple timepoints, clone embeddings reveal dynamics toward common transcriptionally mediated phenotypes—such as activation, cytolytic function or exhaustion—in a more straightforward manner (Figure 1C). The clone embedding can be visualized via UMAP and clustering within this embedding yields the final clonotype profiles. Profile assignments are then projected back onto individual cells, allowing each cell to be probabilistically assigned to its clonotype profile. To clearly differentiate clone embeddings from cell embeddings, all clone embeddings will be shown with a box frame, and all cell embeddings without the frame, in this paper.

Clone embeddings and clone profiles enable a range of downstream analyses that leverage both transcriptomic and clonal information. One application is pseudotime inference (Figure 1D). Conventional pseudotime methods assume that transcriptional distance reflects pseudotime difference and cell transitions occur smoothly within the transcriptomic embedding^7–9, 11, 12^. These assumptions often break down under perturbations, such as drug treatment, where sharp transcriptional shifts or reversion to baseline states can obscure the true developmental ordering within the transcriptomic embedding. Clonotrace addresses this by inferring pseudotime over the clone embedding instead, harnessing shifts in clonal distribution. This clone-informed pseudotime is then mapped back onto individual cells, producing a trajectory that is more robust to discontinuities and state reversals and better aligned with the true temporal structure of cellular differentiation or evolution.

Clonotype profiles also allow the dissection of cell fate decisions. Because they partition cells based on clonal embedding rather than transcriptomic embedding, they can reveal functional differences that would be difficult to detect through gene expression alone. For example, at the root of a differentiating hierarchy—cells may appear homogeneous in expression, yet belong to different clonotype profiles (Figure 1E). Clonotrace uses this information to identify putative fate-driving genes: at the root of a branching trajectory, we compare gene expressions between diverging clonotype profiles after matching cells by pseudotime. By accounting for differences in pseudotime and testing for residual expression differences between profiles, Clonotrace isolates early markers predictive of terminal fate. Even outside of classical differentiation contexts, comparisons between clonotype profiles within a transcriptionally homogeneous cell cluster can reveal biologically meaningful stratification not apparent from the transcriptome alone. In this sense, we view each cell as carrying two complementary identity labels: a static transcriptomic state, and a dynamic clonotype profile. Together, these dual “ID cards” provide a more complete view of a cell’s phenotype at time of sampling as well as its clonal evolution.

### Evaluating Clonotrace on synthetic lineage-tracing data

We first evaluated the capabilities of Clonotrace using two simulated datasets, each generated with real single-cell RNA-seq overlayed with simulated clone labels reflecting distinct differentiation dynamics.

In the first simulation, we assessed Clonotrace’s ability to distinguish between unipotent and bipotent trajectories, a challenge that, to our knowledge, no existing method currently addresses. We used data from a canonical hematopoiesis study comprising 34,782 cells sampled from a branched differentiation process leading from undifferentiated progenitors to monocytes and neutrophils^6^. Cells were clustered into 9 transcriptional states (Figure 2A left). We simulated clones to reflect the true underlying differentiation process that have two terminal fates, monocytes and neutrophils (See Methods for details of the simulation). We defined two clone profiles: a bipotent profile, in which clones give rise to both monocytes and neutrophils, and a unipotent profile, in which clones exclusively differentiate into monocytes. Note that this simulation model may not reflect the true process of hematopoiesis, and is simply for the purpose of method testing. The divergence between the bipotent and mono-potent trajectories was positioned at cell cluster 4 (Figure 2A middle). A total of 3,486 clones were simulated, with 2,370 in the bipotent profile and 1,116 in the mono-potent profile. We allowed asynchronous differentiation speeds between the two branches: for example, the neutrophil lineage was programmed to progress more slowly than the monocyte lineage, as reflected by differences in their time ranges (Figure 2A right). Notably, in the bipotent profile, cells in the neutrophil branch reach the terminal state only after the monocyte lineage has terminated (Figure 2B).

**Figure 2:**
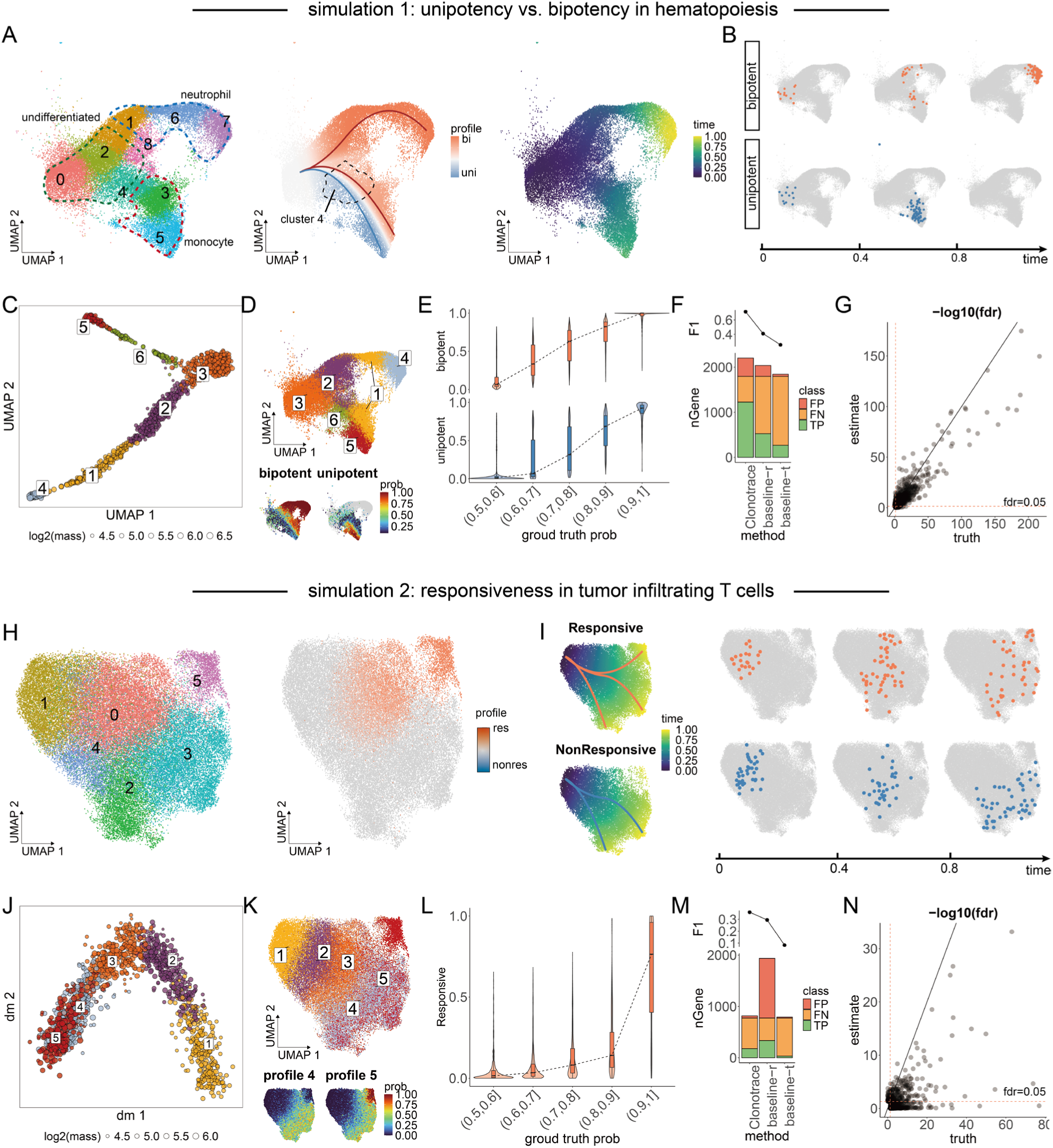
Evaluations on synthetic lineage-tracing data. **A** UMAP of the transcriptomics embedding for the hematopoiesis data. Left, each cell is colored by cluster identity. The regions of three cell types are highlighted. Middle, each cell is colored by simulated probability of clonotype profile assignment. The bipotent profile is marked with red line and the unipotent profile is marked with blue line. Right, each cell is colored by the simulated time. **B** Cell distributions of example clones from the bipotent and unipotent profiles at different time points. **C** UMAP of the clone embedding. Each point is a clone colored by the inferred clonotype profile identity. **D** The same UMAP as **A**, in the top plot each cell is colored by imputed profile identity. In the bottom plots, each cell is colored by the summed probability of being assigned to bipotent-associated profiles (1,2,4; left) or unipotent-associated profiles (5, 6; right). **E** Comparison of inferred versus ground-truth profile probabilities for bipotent and unipotent lineages. **F** Comparison of fate-driving gene detection performance and F1 score between Clonotrace and baseline methods (TP: true positive; FP: false positive; FN: false negative). **G** Comparison of FDR values for significant genes identified using ground-truth versus Clonotrace-inferred profile probabilities. **H** UMAP of cell transcriptomics embedding for infiltrating T cell data. Left, Cells colored by cluster identity. Right, Cells colored by simulated probability of assignment to the responsive profile. **I** Left: the trajectory for the responsive clones (top) and non-responsive clones (bottom), each cell is colored by the simulated time. Right: the cell distribution in the responsive and nonresponsive clones at different time points. **J** The diffusion map of the clone embedding. Each point is a clone colored by the profile identity. **K** The same UMAP as **H**. In the top plot, each cell is colored by imputed profile identity. In the bottom plot, each cell is colored by the probability to be assigned into profile 4 (left) or 5 (right). **L** Comparison of inferred versus ground-truth profile probabilities for responsive profile. **M-N** Same analysis as in panels **F-G**, applied to the T cell simulation.

We applied Clonotrace to the 1,210 expanded clones (with ≥10 cells; Supplementary Figure 3A). The resulting clone embedding revealed clear bifurcation, and clustering with an arbitrary cluster number of six gives clonotype profiles as shown in Figure 2C, left. When projected back onto the transcriptomic embedding, profile 3 captured the undifferentiated population, profiles 1, 2, and 4 captured successive stages along the bipotent trajectory, and profiles 5 and 6 corresponded to the unipotent lineage (Figure 2C right, Figure 2D). Due to the slower speed of neutrophil differentiation, bipotent clones that ultimately produce only neutrophils were grouped into profile 4. Clonotrace-inferred probabilities of unipotency (profiles 5 and 6) versus bipotency (profiles 1, 2, 4) were strongly correlated with their ground-truth probabilities (Pearson’s r = 0.890 for the bipotent lineage, r = 0.794 for the unipotent lineage; Figure 2E). We observed a modest underestimation of profile probabilities, particularly for undifferentiated cells with weak fate bias (true probabilities near 0.5), which were primarily assigned to the intermediate profile 3 that contains mostly cells prior to fates determination.

To identify genes associated with the early fate bias between unipotency and bipotency, we performed differential expression analysis between profile 1 (bipotent) and profile 6 (unipotent) within cell cluster 4, the point of their divergence. As ground truth, we used the linear regression model (Supplementary Figure 3E) to identify fate-driving genes based on the simulated “true” clone profile labels and simulated “true” time, yielding 1,898 significant genes after FDR correction. Clonotrace applies the same linear regression model, but with the inferred profile probabilities and inferred pseudotimes (Supplementary Figure 3B-D) in place of the ground-truth annotations as input. As there are no other methods for detecting bipotency versus unipotency associated genes in the absence of temporal information, we benchmarked against simpler “baseline” methods which proceeded as follows: First, cells were classified as bipotent progenitors if their clones extended to terminal clusters 1, 6, or 7, and as unipotent progenitors otherwise (Supplementary Figure 3F). This uses the knowledge, masked to Clonotrace, that bipotent cells differentiate towards these clusters, while unipotent cells do not. Differential expression (DE) testing between these two groups in cluster 4 revealed putative fate-associated genes. However, because the bipotent and unipotent groups differed in their time distributions, this comparison conflates fate bias with developmental stage. To mitigate this, we performed a time-matched version of the same analysis, selecting subsets of cells from both groups with similar time distributions (Supplementary Figure 3G). These baseline comparisons — both raw (baseline-r) and time-matched (baseline-t) — served as references for comparison. Clonotrace recovered 1,199 of the 1,898 ground-truth genes (63%) with a false positive rate of 11% (154 genes). Compared to both baseline DE tests, with or without time matching, Clonotrace achieved the highest F1 score (0.713) in identifying fate-driving genes (Figure 2F). This indicates that, compared to the baseline method, Clonotrace’s profile assignment and profile-level DE improves power and robustness. A comparison of FDR values between the ground-truth and Clonotrace-inferred results showed that most false positives and false negatives occurred near the statistical significance threshold (Figure 2G), indicating robust recovery of core bifurcation-associated genes.

While the hematopoiesis simulation featured a clearly bifurcating trajectory visible in the transcriptomic embedding, our second simulation was designed to mimic T cell dynamics in disease contexts where cell states are less distinct, and differentiation signals are more nuanced. We used a single-cell RNA-seq dataset from tumor-infiltrating CD8+ T cells in glioblastoma, which we will analyze in detail later in this paper. 46801 cells were clustered into six subsets based on gene expression (Figure 2H left). Then, we simulated a differentiation process beginning at cluster 1 and branching into three terminal states: clusters 2, 3, and 5. To mimic the scenario where, upon treatment, certain T cell clones diversify into new fates, we simulated 2813 clones in two profiles: non-responsive clones that follow a differentiation trajectory from cluster 1 through clusters 0 and 4 to terminal states 2 and 3, and responsive clones that additionally exhibit increased probability of differentiating from cluster 0 into cluster 5 (Figure 2H right, I). Importantly, this probability forms a gradient across cells in cluster 0, reflecting the subtle fate bias.

Compared to the hematopoiesis simulation, this dataset presents a greater challenge for both clone profile inference and gene identification, due to the probabilistic nature of the responsive fate where only a small subset of cells split into the divergent state (cluster 5). Difficulty is also increased by the large overlap of intermediate states between clonotype profiles. Applying Clonotrace to the 1399 expanded clones (Supplementary Figure 3H), we identified five clonotype profiles (Figure 2J). Profiles 1 to 3 captured the shared distribution from root-to-intermediate states common to both responsive and non-responsive clones, while profiles 4 and 5 correctly distinguished the distinct terminal fates (Figure 2K). Cluster 0 contained cells from both profiles and was transcriptionally most similar to cluster 5, which was exclusive to the responsive profile.

Clonotrace-inferred probabilities of responsive profile (profiles 5) were positively correlated with the ground-truth probabilities (Pearson’s r = 0.664) (Figure 2L). To identify early predictors of responsiveness, we performed DE analysis within cluster 0, comparing cells assigned to the responsive profile (profile 5) against those in other profiles. Using the same benchmark approach as in the hematopoiesis simulation (Supplementary Figure 3I-N), we identified 776 significant fate-driving genes using ground-truth labels. Clonotrace identified 228 fate-driving genes, of which 183 were true positives—highlighting the inherent difficulty of detecting subtle, probabilistic fate biases in noisy single-cell data. In contrast, while the baseline-r method found slightly more genes, it suffers from inflation of false positives. The baseline-t method, due to pseudotime matching, had good specificity but almost no power. Overall, Clonotrace achieved the highest F1 score (0.365) compared to baseline methods (Figure 2M), successfully differentiating the responsive profile 5 from the unresponsive profile 4 and recovering the strongest lineage-driving signals (Figure 2N). This example shows that Clonotrace can yield informative results in an extremely challenging scenario.

### Clonotrace reconstructs temporal order across abrupt transcriptomic shifts

Traditional pseudotime inference methods assume that transcriptional distance reflects pseudotime difference and that cell transitions are smooth within the transcriptomic embedding. However, this assumption can break down if intermediate transitional states are missing from the data, or if cells follow cyclic or complex trajectories that reverse course. To test whether Clonotrace can recover correct temporal ordering of cells under conditions that violate the assumptions, we analyzed a lineage-traced cancer cell line dataset involving three distinct treatments (Figure 3A). This dataset^32^ was generated using a lentiviral barcoding system that stably labels each cell and its descendants with a unique DNA barcode. After barcode integration and brief expansion, the cells were split into three groups, each subjected to one of the treatments: CoCl₂ to mimic hypoxia, a combination of dabrafenib and trametinib as targeted melanoma therapy, or cisplatin as a DNA-damaging chemotherapeutic. Cells were collected at three key time points: day 0, day 10 (short-term response), and week 5 (long-term adaptation). We used the single-cell RNA-seq and barcode sequencing data from each timepoint.

**Figure 3:**
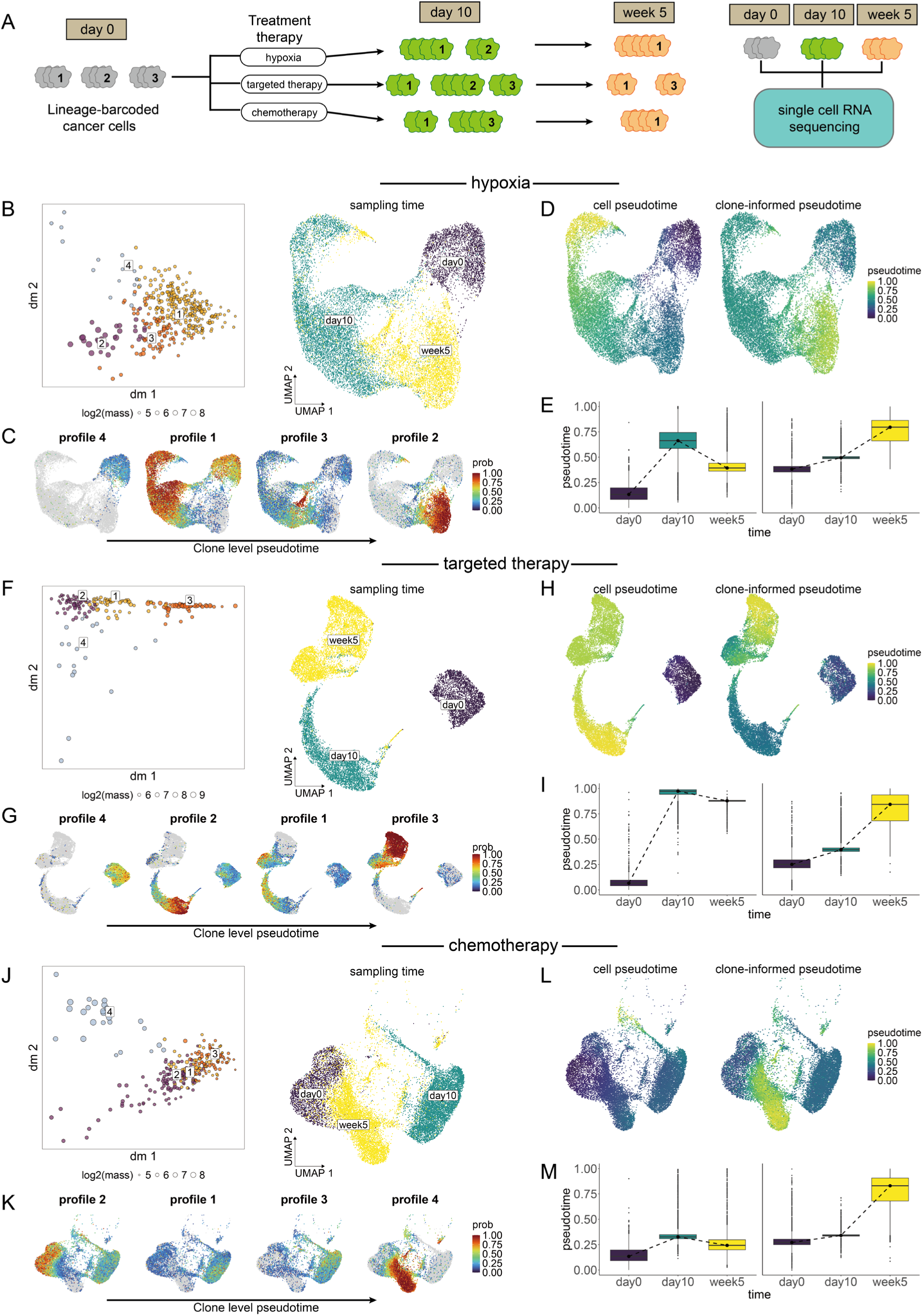
Clonotrace reconstructs temporal order across abrupt transcriptomic shifts. **A** Schematic of the treatment experiment. **B** Left, diffusion map of the clone embedding. Each point is a clone colored by profile identity. Right, UMAP of cells treated using hypoxia, colored by sampling time. **C** Probability density of each clone profile projected onto the cell UMAP. Profiles are ordered by clone-level pseudotime. **D** Comparison of pseudotime derived from cell embedding (left) and clone embedding (right), shown over the cell UMAP. **E** Boxplot comparison of cell-level pseudotime and clone-informed pseudotime. **F-I** Corresponding panels for cells treated with targeted therapy, aligned in the same layout as panels B–E. and **J-M** Corresponding panels for cells treated with chemotherapy, aligned in the same layout.

This system provides a rigorous test case for Clonotrace: under all treatments, cells exhibit abrupt transcriptional changes, with few or no transitioning cells captured between day 0 and day 10. For example, in the hypoxia condition, transcriptomic profiles at day 10 are highly divergent from baseline, reflecting acute stress responses. By week 5, however, cells begin to return toward a pre-treatment–like expression state, creating a *boomerang-like effect* in the transcriptomic space (Figure 3B right). When diffusion pseudotime^11^ method is applied to the cell embedding using day 0 as the root, the algorithm incorrectly positions week 5 between day 0 and day 10 due to its reliance on transcriptional similarity rather than biological time (Figure 3D, E). This failure arises because traditional pseudotime inference is based on proximity in the transcriptomic manifold—an assumption violated here due to treatment-induced bottlenecks and missing intermediate (transitioning) cells.

In contrast, Clonotrace leverages the temporal continuity of clonal behavior rather than transcriptomic proximity. Each clone’s embedding is based on the distribution of its member cells in transcriptomic space. Changes in this distribution can be continuous, even when changes of its member cells are abrupt and discontinuous. When applied to the same hypoxia dataset, Clonotrace identified four clone profiles that together form a smooth progression in clone space(Figure 3B left). Using the clone most enriched in day0 as the starting point, we derived the clone-level pseudotime via diffusion on the clone embedding. The clone-level pseudotime time ordered the profiles as 4->1->3->2 (Supplementary Figure 4). When these profiles are projected back onto the cell embedding and arranged by the clone-level pseudotime, we observed that: profile 4 is enriched at day 0, profile 1 dominates at day 10, and profiles 2 and 3 emerge at week 5 (Figure 3C), accurately reconstructing the experimental timeline. The clone-level pseudotime is further smoothed across the cell embedding, faithfully capturing the true temporal progression while preserving continuity in cell state transitions. (Figure 3D, E).

Similar results were observed in the targeted therapy and chemotherapy treatment arms. In both cases, Clonotrace’s clone-informed pseudotime aligned well with experimental time points, even when the cells formed isolated clusters whose proximity in gene expression embedding do not agree with their true temporal progression. The clone embedding effectively connected transcriptionally disjoint cell states (Figures 3F–I, 3J–M) revealing trajectories that conventional methods fail to resolve.

### Clonotrace distinguishes unipotent versus bipotent cells in hematopoietic differentiation

We next applied Clonotrace to a public lineage-barcoded dataset of in vitro mouse hematopoiesis^6^ to evaluate its performance in a well-characterized but biologically complex system. Hematopoietic stem and progenitor cells were labeled with genetically heritable barcodes at day 0 and cultured until day 6. Cells were sampled for single-cell RNA sequencing at days 2, 4, and 6 (Figure 4A). At each time point, clones were defined as groups of cells sharing the same lineage barcode. Across timepoints, Clonotrace was used to construct a clone embedding, where the same clone across different timepoints is given separate embedding values. The embedding was then used to cluster the clones into 8 distinct profiles (Figure 4B left). In the clone embedding, there is clear fan-shaped geometry with profile 8 at one tip. Profile 8 contains exclusively hematopoietic progenitor cells, and thus we can assume that it is the initiating profile. Retrospective linking of the same clones across days 4 to 6 confirmed the transitions from profile 3 to profile 7, from profile 5 to profile 6, and from profile 1 to profile 4 (Figure 4B right). To better visualize the layout of clone profiles within the cell embedding, we introduce clone-weighted cell embedding. This embedding was constructed by reweighting edges in the cell-cell nearest-neighbor graph—originally defined based on transcriptomic similarity— to incorporate the neighborhood relationships of their associated clones in the clone embedding (see Methods for details). In the cell embedding, based on marker expression, we identify the trunk containing progenitor cells leading to the monocyte and neutrophil states. After we projected the profile identities on to the clone-weighted cell embedding, we observed that the transition between profile 3 to 7 corresponds to neutrophil differentiation, while the transition from profile 5 to 6 corresponds to monocyte differentiation. In addition to these mono-potent trajectories, the transition from profile 1 to profile 4 corresponds to bipotent differentiation towards both monocyte and neutrophil fates (Figure 4C left). Three different clones, representing each of unipotent monocyte differentiation, unipotent neutrophil differentiation, and bipotent differentiation, are shown in Figure 4D. Notably, in clone-weighted embedding, unipotent and bipotent monocytes are clearly separated, despite being indistinguishable in transcriptomic space (Figure 4C, right). The separation between unipotent and bipotent monocytes corroborates the distinction between neutrophil-like and dendritic-like monocytes previously described by Weinreb et al. A similar separation was also observed for neutrophils within cell clusters 6 and 7.

**Figure 4:**
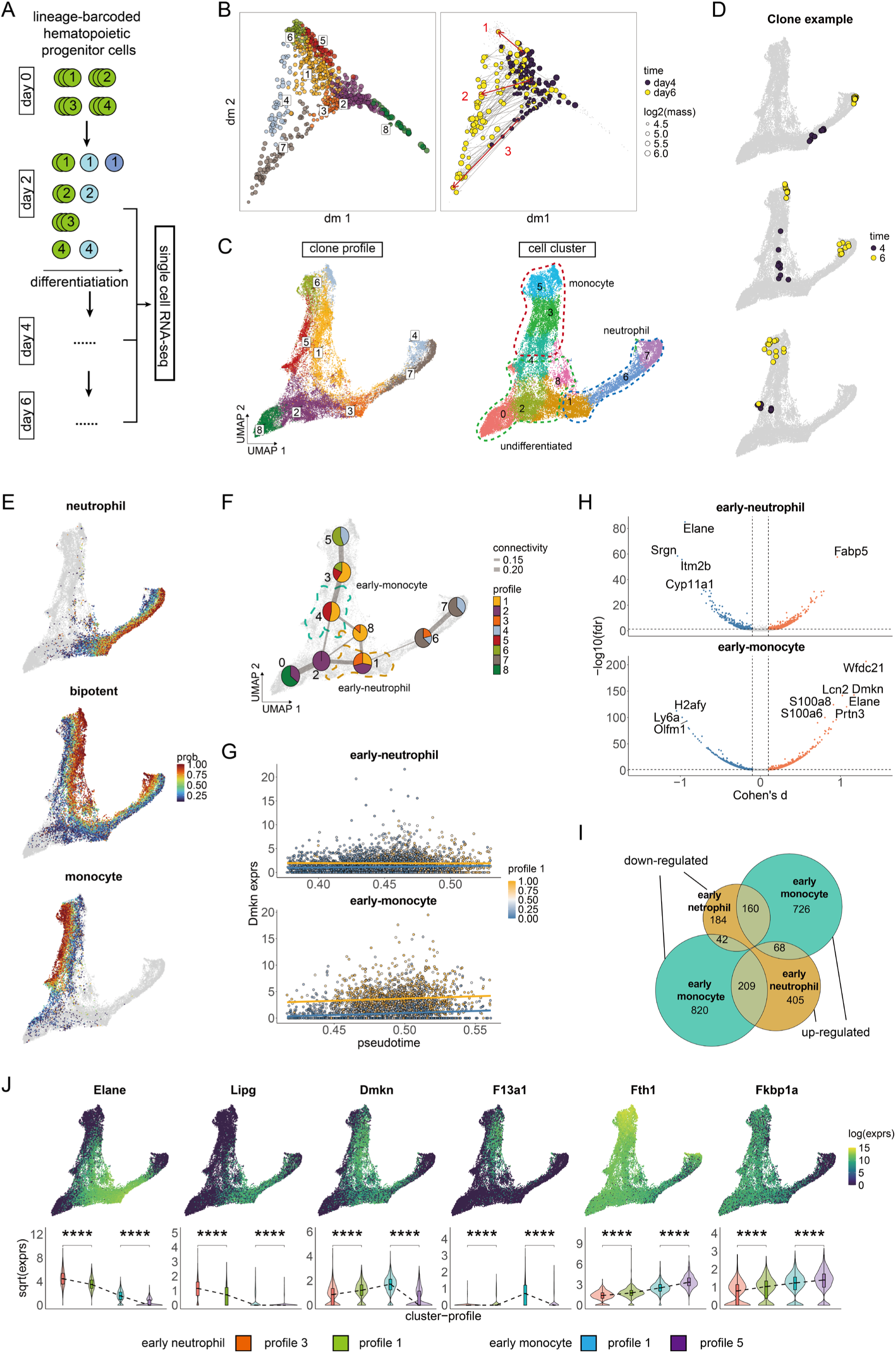
Clonotrace distinguishes unipotent from bipotent cells in hematopoiesis. **A** The scheme of the lineage-barcoded experiment for hematopoiesis in mouse. Each circle is a cell colored by differentiation stage and labeled by the lineage barcode. **B** The diffusion map of the clone embedding. Left, each point is a clone colored by profile identity. The center of each profile is marked with the profile label. Right, clones which are detected in both day 4 and day 6 are highlighted using steel blue and coral and linked by an arrow. Three example clones which represent different profile transitions are highlighted with red circles and arrows. **C** The UMAP of clone weighted cell embedding, each point is a cell. Left, cells colored by the profile identity (probability larger than 50% for the corresponding profile). The center for each profile is marked. Right, cells colored by cluster identity. **D** The cell distributions for the three example clones highlighted in **B** right, labeled with corresponding id. Cells within the target clone are colored by time points. **E** The cell distribution for the mono-potent neutrophil, bipotent and mono-potent monocyte profiles. Each cell is colored by the probability to be classified into this profile (cells with probability lower than 0.1 are colored in grey). **F** The scatter pie plot shows the proportion of profiles within each cluster. Each node is a cluster colored by the composition of profiles. Nodes are linked by edges generated from PAGA^9^. The cluster 1 and 4 in which the fate divergence happened are highlighted with dashed contour (1: yellow; 4: green). **G** The example to show the DEG detection by linear regression for profile 1 in cluster 1 and 4. Each point is a cell colored by the probability to be assigned to profile 1. Regression lines for profile 1 (yellow) and other profiles (blue) are overlaid. **H** The volcano plot to show the significant genes detected for the bipotent profile 1 in cluster 1 (upper) and cluster 4 (bottom). **I** The Venn diagram to show the overlap between significant genes for the bipotent profile 1 in cluster 1 and cluster 4. **J** Top, the log-wise expression for example genes over the cell UMAP. Bottom, the comparison of gene expression (transformed by square-root) between the three profiles in cluster 1 and 4.

For each cell, we mapped the probability of unipotency (neutrophil or monocyte) versus bipotency (Figure 4E). To determine if there are early transcriptional markers of bipotent versus unipotent differentiation, we first performed an enrichment test to detect cell clusters enriched for cells from the bipotent profile 1 (Supplementary Figure 5A) and then conducted differential expression analysis between profile 1 and other profiles within this cluster. The earliest transcriptional boundaries appeared in cluster 4 (between profiles 1 and 5) and cluster 1 (between profiles 1, 2, and 3) (Figure 4F). Clusters 1 and 4 marked the beginning of differentiation into, respectively, monocytes and neutrophils. Thus, we will call cluster 1 “early neutrophil” and cluster 4 “early monocyte”. Using our linear regression–based method adjusting for pseudotime (Figure 1E, Supplementary Figure 5 B-D), we compared gene expression between profile 1 and other profiles within these two clusters (Figure 4G). In total, we identified 1,121 significant genes in early neutrophils and 2,098 significant genes in early monocytes for profile 1 (FDR < 0.05) (Figure 4H, Supplementary Data 1). Among genes with appreciable effect sizes (absolute Cohen’s d > 0.1), 954 were upregulated in early monocytes for the bipotent profile. Of these, 160 were significantly downregulated in early neutrophils (Figure 4I, Supplementary Figure 4E), including canonical neutrophil markers such as *Elane*, *Mpo*, and *S100a8/a9* (Supplementary Figure 4F). UMAP visualizations of gene expression for *Elane* and other representative genes (Figure 4J) show high expression in the neutrophil branch and low expression in the monocyte branch, while cells belonging to the bipotent profile—located in between—exhibit intermediate expression levels. This gradient is also evident in the corresponding violin plots, which show a clear decrease in expression from neutrophil to bipotent to monocyte cells. Conversely, in early neutrophils, 209 out of 682 bipotent-upregulated genes were downregulated in early monocytes (Figure 4J right). Only 68 genes are bipotent-upregulated in both early monocytes and early neutrophils (Figure 4J middle). These findings suggest that bipotent differentiation is mainly characterized by intermediate expression of gene programs associated with both neutrophil and monocyte lineages, distinguishing it transcriptionally from unipotent trajectories.

### Clonotype profiles uncover recurrent, fate-divergent CD8⁺ T cell programs in glioblastoma

To demonstrate how Clonotrace can be used to interrogate T cell dynamics, we analyzed a dataset of tumor-infiltrating CD8⁺ T cells from 44 glioblastoma patients spanning a range of tumor grades and treatment conditions (Figure 5A)^33^. The dataset comprises 129,465 CD8⁺ T cells profiled by joint scRNA-seq and TCR-seq. Using the CDR3α/β amino acid sequence, we assigned cells to clones within each patient and applied Clonotrace to identify clonotype profiles that capture shared patterns of clonal distribution in gene expression space.

**Figure 5:**
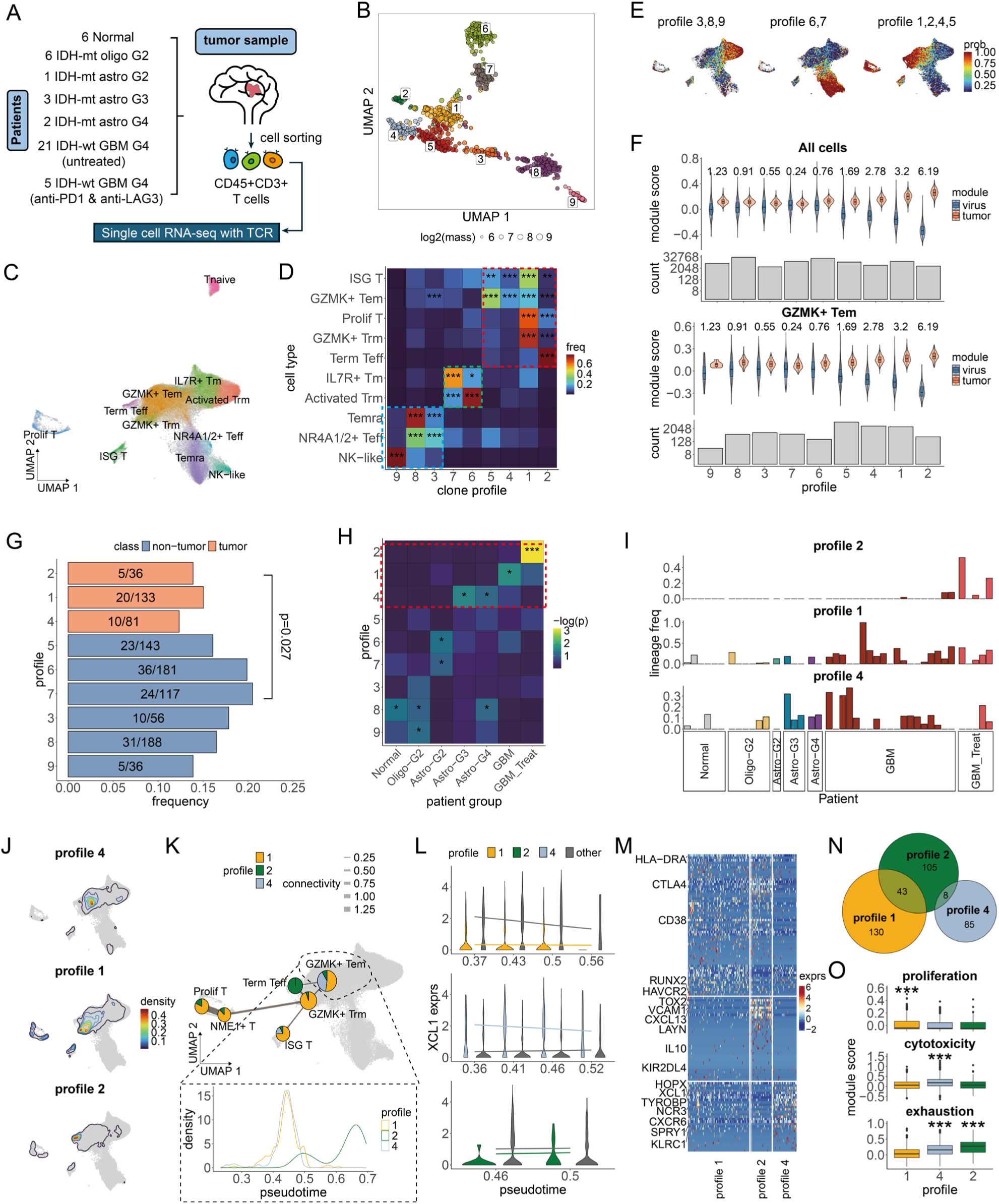
Analysis of CD8 T cells in tumor micro-environment in glioblastoma. **A** Overview of patients’ sample collection and cell sequencing. **B** UMAP of cell transcriptomic embedding, each point is a cell colored by cell type. **C** UMAP of clone embedding, each point is a clone colored by clone profile. **D** Cell type enrichment of each profile. The color of each tile indicates the frequency of cells for the given profile in the given cell type (normalized within each cell type). Significant enrichments are highlighted with *. **E** Cell probability density of the three profile groups delineated in D on the cell UMAP. **F** Comparison of the module score for tumor-reactive vs. virus-reactive genes in each profile across all cells (top) and within GZMK+ Tem cells (bottom). Cohen’s d between tumor and virus-reactive module scores are shown at the top. Cell count for each profile is shown in the bar plot at the bottom. **G** The bar plot shows the frequency of clones annotated with a virus antigen. The number of annotated clones and total clones are marked for each profile with the format annotated/total. Tumor reactive profiles are highlighted in coral. **H** Profile enrichment in different tumor grades. Each tile is colored by the log scaled p-value for the enrichment test (one-side Wilcox rank sum test). Significant enrichments are highlighted with *. **I** Clone distribution in different patients across tumor grades in tumor reactive profiles. Patients are ordered and colored by tumor grade. The height of the bar denotes the frequency of clones in the given profile for the patient. **J** Contour plot to show the cell probability density for profiles 1, 2 and 4. **K** The top plot shows the PAGA network for cell types enriched in profile 1, 2 and 4. Each node is a cell type colored by the profile composition. The GZMK+ Tem population is highlighted with a black dashed circle. The bottom plot shows the cell density distribution along pseudotime separately for profiles 1, 2, 4 in GZMK+ Tem. **L** An example to illustrate DEG detection by linear regression on pseudotime for profiles 1, 2, 4 within GZMK+ Tem cells. The violin plot shows the expression distribution for cells in the target profile and other profiles in a pseudotime range. Regression lines for the target profiles (1: yellow; 2: green; 4: blue) and other profiles (grey) are overlaid. **M** Heatmap to show the expression of upregulated genes in the GZMK+ Tem cells in each profile, with genes ordered based on their effect size, profile 1 genes are shown first, followed by profiles 2 and 4. Selected marker genes are shown in the row names. **N** Venn diagram to show the number of genes which are significantly upregulated in each profile in the GZMK+ Tem cells. **O** Comparison of the module score between profiles within GZMK+ Tem cells. Profiles with significantly high module scores are highlighted with *. (* for <0.1; ** for <0.01; *** for < 0.001).

Clonotrace identified 9 clonotype profiles from 971 expanded clones (i.e. clones with ≥10 cells) (Figure 5B, Supplementary Figure 2A). All profiles were shared across patients, suggesting that they reflect recurrent modes of clonal behavior rather than patient-specific effects (Supplementary Figure 6A-B). Unlike standard transcriptomic clustering, which partitions cells into disjoint regions in the transcriptomic embedding (Figure 5C), each clonotype profile mapped onto distinct regions of the transcriptomic landscape but did so in a non-exclusive manner (Figure 5D). For instance, profiles 3, 8, and 9 were enriched in Temra, NK-like, and NR4A2⁺ effector subsets; profiles 6 and 7 were enriched in IL7R⁺ memory and Trm cells; while profiles 1, 2, 4 and 5 were enriched in GZMK⁺ Tem, ISG⁺, and terminal effector (Teff) populations (Figure 5E). This shows that while transcriptional states may overlap between T cell clones, they are not functionally equivalent: cells with similar transcriptomes often belonged to clonotypes with different profile identities, suggesting differing underlying dynamics.

First, we identified tumor-reactive clonal profiles using published gene modules that distinguish tumor-from virus-specific CD8⁺ T cells (Supplementary Data 2). Profiles 1, 2, 4 and 5 showed significantly higher tumor-reactive module scores relative to virus-reactive modules (Figure 5F, top), implicating them as putative tumor-reactive clonotype profiles. Crucially, this pattern persists after stratification by transcriptomic state. For example, among GZMK⁺ Tem cells, which have modest associations with tumor-reactivity (Supplementary Figure 6C), profiles 1, 2, 4 and 5 retained significantly higher tumor-reactivity scores than other profiles (Figure 5F, bottom). This underscores that clonotype profiles, when layered on to transcriptome-based stratification, can reveal additional functional heterogeneity.

To assess whether the tumor-reactive clonotype profiles identified by Clonotrace align with antigen specificity, we examined antigen annotations for 167 CDR3β sequences with known viral specificities. While most public annotations are biased toward viral antigens due to limited tumor antigen data, we found that profiles 1, 2, and 4 exhibited lower frequencies of virus-specific TCRs, with borderline statistical significance (p = 0.027, binomial test; Figure 5G). Although the magnitude of this depletion was modest, the trend is consistent with the hypothesis that these clonotype profiles are enriched for tumor-reactive clones. At the same time, the weak separation based on TCR sequence annotations highlights a critical limitation of current TCR sequence– based approaches: in heterogeneous disease settings like cancer, where antigenic landscapes are diverse and often patient-specific, sequence similarity alone is often insufficient to stratify T cell function or fate.

To further underscore the difference with TCR sequence-based approaches for clone stratification, we performed unsupervised clustering of clones based on TCRdist3^34^, a method that defines clonal distances via sequence similarity of TCR αβ sequences. We found that CDR3β sequence distances had minimal correlation with transcriptomic distances between clones (Spearman’s ρ = 0.022; Supplementary Figure 6D–E). While clones with nearly identical CDR3β sequences sometimes shared similar distributions in transcriptomic space, this relationship deteriorated rapidly as sequence similarity decreased (Supplementary Figure 6F). Moreover, the TCRdist-based clusters showed poor alignment with transcriptional structure and failed to separate tumor-reactive from virus-reactive populations—contrasting sharply with the structure uncovered by Clonotrace (Supplementary Figure 6G–H). This contrast emphasizes that while TCR sequence can provide valuable information in controlled antigen contexts, it lacks the resolution to distinguish functional states or differentiation trajectories in complex, polyclonal environments like cancer.

Importantly, clonotype profiles exhibited coarse-grained clinico-genomic stratification (Figure 5H). Profiles 1, 2, and 4—those with highest tumor-reactive scores—were significantly enriched in patients with high-grade gliomas (e.g., Astro-MUT-G4, GBM-WT), regardless of treatment (Figure 5I), suggesting they reflect immunological states associated with aggressive disease. These patterns were not driven by sampling or batch effects: all profiles included clones from multiple patients (Supplementary Figure 6B), and no profile was restricted to a specific patient group.

Thus, we focused on the tumor-reactive profiles 1, 2, and 4, all three of which converge in a GZMK⁺ Tem subset but extended to distinct terminal fates (Figure 5J). Profile 1 progressed toward a GZMK⁺ Trm fate; profile 2 extended to both GZMK⁺ Trm and terminal Teff, with the highest density in terminal Teff; and profile 4 progressed towards Temra. PAGA analysis further confirmed that the GZMK⁺ Tem subset is centrally situated in the transcriptomic embedding (Figure 5K, top), suggesting these clonotypes begin in a shared, fate-flexible state but diverge transcriptionally along distinct developmental axes. Pseudotime analysis of cells within the GZMK⁺ Tem subset indicated that profiles 1 and 4 are at a similar stage of differentiation, while profile 2 represents a more advanced state (Figure 5K, bottom, Supplementary Figure 6I-J).

To identify genes associated with fate divergence, we performed differential expression within GZMK⁺ Tem cells across the three profiles. For each profile, we compared its cells against the other cells within the GZMK⁺ Tem subset, after stratification by pseudotime (an example for *XCL1* is shown in Figure 5L). In total, we identified 173 genes upregulated in profile 1, 156 genes upregulated in profile 2, and 93 genes upregulated in profile 4(FDR < 0.05, Cohen’s d > 0.1; Supplementary Data 3). Profiles 1 and 2 shared the upregulation of *RUNX2*, a Trm-associated transcription factor, as well as proliferation (*MKI67*, *TYMS*), activation (*CD38*), and inhibitory (*CTLA4*) genes. This is consistent with a cycling, transitional phenotype. Profile 2 alone upregulated exhaustion markers including *TOX2* and *CXCL13*, while profile 1 upregulated cell-cycle regulators (*CDK1*, *TOP2A*), indicating divergence between proliferative and exhausted branches (Figure 5M–O). In contrast, profile 4 showed upregulation of cytotoxicity genes (*XCL1*, *NCR3*), consistent with direct progression toward the Temra state.

These findings illustrate how Clonotrace allows the identification of shared and clinically relevant patterns of T cell dynamics across patients. In this cancer setting, Clonotrace reveals that tumor-reactive clonotypes converge onto a shared, fate-flexible GZMK⁺ Tem state that is connected to three distinct outcomes: proliferative Trm-like, cytotoxic effector, and exhausted. These trajectories are not readily apparent from transcriptional similarity alone but emerge through clonotype profiling. Clonotrace further delineated fate boundaries that are not evident through transcriptional profiling, enabling the identification of putative fate driving genes. Please see Wang and Fu et al.^33^ for a more extensive analysis of this data, including a discussion of the clinical significance of these results.

### Clonotrace reveals longitudinal dynamics of circulating CD8⁺ T cells during immunotherapy

To demonstrate how Clonotrace can resolve complex T cell dynamics in a longitudinal setting, we analyzed scRNA-seq and TCR-seq data from peripheral blood CD8⁺ T cells in patients with non–small cell lung cancer (NSCLC). Eight patients were enrolled in a treatment protocol that began with two cycles of anti-PD1 monotherapy, followed by two cycles of anti-PD1 combined with a JAK1 inhibitor (JAKi) and then continued with only standard of care anti-PD1 (Figure 6A). A subset of patients who initially failed to respond to anti-PD-1 monotherapy demonstrated clinical responses when treated with the combination of anti-PD-1 and a JAK inhibitor, consistent with findings from preclinical studies and clinical trials in Hodgkin lymphoma^35^. We reanalyzed this dataset using Clonotrace to investigate clonal dynamics throughout treatment and identify patterns associated with clinical response.

**Figure 6:**
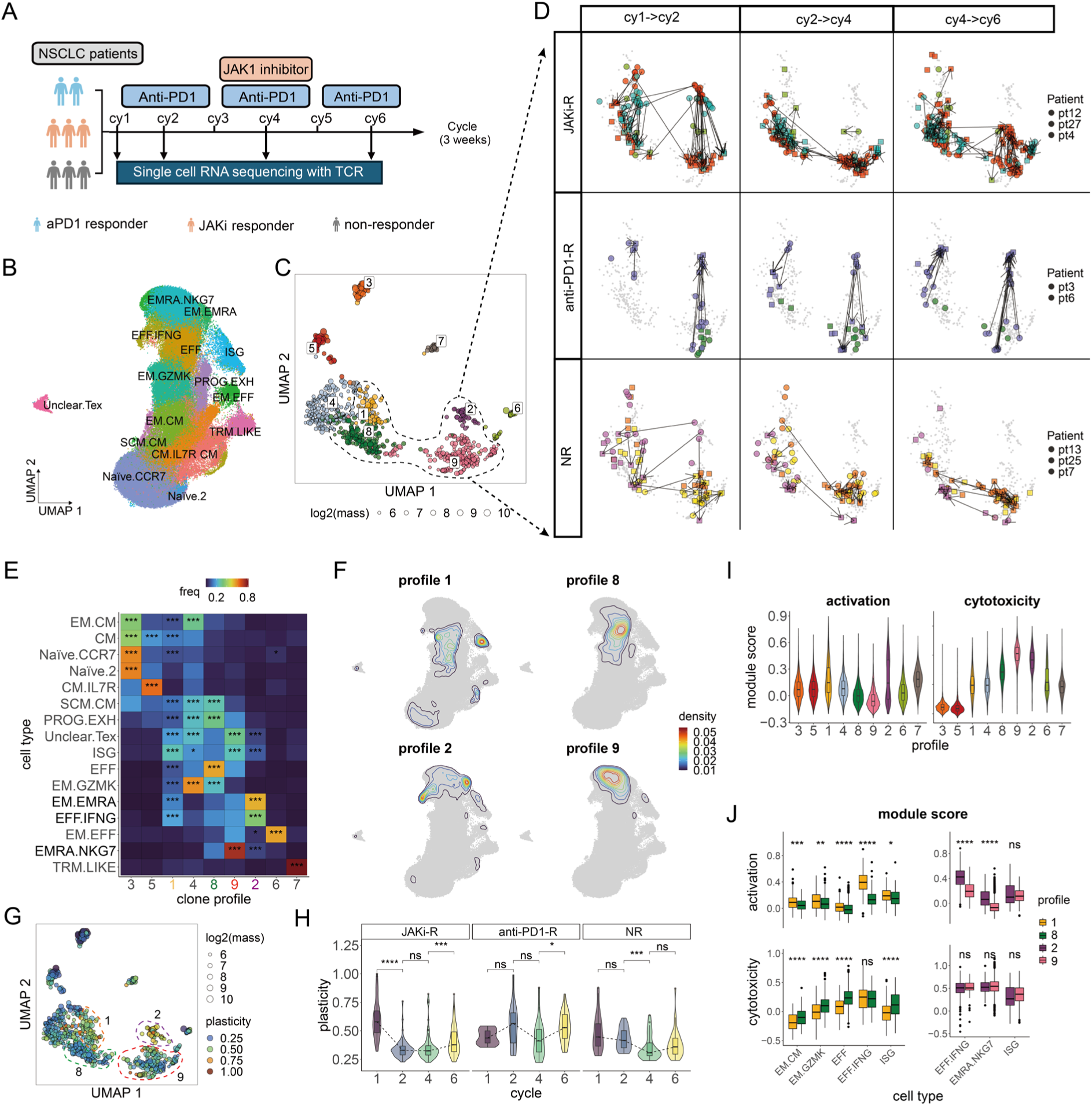
Analysis of longitudinally sampled CD8 T cells in PBMC from cancer patients in immunotherapy. **A** The scheme of the treatment procedure. **B** The UMAP of cell transcriptomic embedding, each point is a cell colored by cell type. **C** The UMAP of the clone embedding, each point is a clone colored by clone profile. The size of the point denotes the log transformed cell amount in that clone. The focused profiles 1, 8, 2 and 9 are highlighted with a dashed circle. **D** The clone transition during the treatment. All plots are the zoom-in version of clone UMAP in **C** for profile 1, 2, 8 and 9, each row is a response group, and each column is a treatment cycle. Each point is a clone colored by the patient source. Clones with the same TCR at different cycles are connected by an arrow. **E** The cell type enrichment in each profile. The color of each tile indicates the frequency of cells for a given profile in the cell type (normalized within a cell type). Significant enrichment is highlighted with *. **F** The cell probability density of the profile 1, 8, 2 and 9 over the cell UMAP. **G** The same clone UMAP as **C**, each point is a clone colored by plasticity. **H** Comparison of clonal plasticity with profile 1, 2, 8 and 9 during treatment in each response group. Significant change in plasticity is highlighted with *. **I** The comparison of the module score for cell activation and cytotoxicity between profiles. **J** The comparison of the module score for cell activation and cytotoxicity between profile 1 and 8, 2 and 9 within each cell type. Significant difference in module score is highlighted with *. (* for <0.1; ** for <0.01; *** for < 0.001).

Patients from three different clinical response groups were analyzed: two anti-PD1 responders, who responded during the initial monotherapy phase (anti-PD1-R); three JAKi responders, who failed to respond during the anti-PD-1 monotherapy phase but subsequently responded when JAKi was added to the anti-PD-1 (JAKi-R); and three non-responders, who failed to respond even after the addition of JAKi (NR). Peripheral blood mononuclear cells (PBMCs) were collected from all patients at cycles 1, 2, 4, and 6, and profiled using scRNA-seq and TCR-seq, yielding a dataset of 150,150 CD8⁺ T cells assigned to 16 transcriptionally defined subsets (Figure 6B). Clones were defined by identical CDR3α and CDR3β amino acid sequences within each patient and timepoint. To capture clonal behavior across time, separate embeddings were learned for clones with the same TCR sampled at different timepoints, and the shift in these embeddings allowed for the dynamic tracking of clone distributions.

Across all patients and timepoints, Clonotrace inferred nine clonotype profiles (Figure 6C), each enriched for distinct CD8⁺ T cell subsets (Figure 6E). These profiles were shared across patients, and thus they reflect reproducible clonal behavior patterns rather than patient-specific artifacts (Supplementary Figure 7A-B).

To dissect treatment-induced T cell dynamics, we overlaid sampling time onto the clone embedding and linked identical clones sampled at adjacent timepoints. We defined a clonal shift as a change in profile identity for a given clone between timepoints. Three intervals were examined: cycle 1 to cycle 2 (anti-PD1 monotherapy), cycle 2 to cycle 4 (addition of JAKi to anti-PD1 therapy), and cycle 4 to cycle 6 (anti-PD1 monotherapy). Comparing clonal shifts across these intervals and across patient response groups revealed pronounced differences between the three patient groups (Supplementary Figure 7C), particularly involving clonal shifts between profiles 2 and 9, and between profiles 1 and 8 (Figure 6D).

It is important to note that “clonal shift” here refers to a change in the density distribution of a clone’s cells within the transcriptomic embedding, not necessarily direct differentiation. Such shifts can reflect clonal expansion, contraction, or migration out of peripheral blood.

First, we focused on the pronounced pattern that emerged between profile 2 and profile 9 (Figure 6D). Throughout the course of treatment, the NR patients have very few clones in profile 2, while the anti-PD1-Rs and JAKi-Rs showed strong but opposite clonal shifts between these two profiles: During the initial anti-PD1 monotherapy (cy 1–>2), one of the two anti-PD1-Rs exhibited a clear clonal shift from profile 9 to 2 involving 6 TCR clones. In contrast, two of the three JAKi-Rs showed a clear shift in the opposite direction from profile 2 to 9 involving 17 TCR clones, despite having no initial clinical response to anti-PD1. This suggests that JAKi-Rs possessed clonotypes that responded to anti-PD1 prior to the introduction of JAKi, even though these patients did not respond clinically to anti-PD1 during this time interval. In contrast, the NRs were static and did not show shared trends that are shared between TCR clones.

Then, upon JAKi administration (cy2->4), the same anti-PD1-Rs reversed trajectory from profile 2 back to 9. Meanwhile, the clones in JAKi-Rs mostly remained in profile 9. Following the withdrawal of JAKi (cy4->6), anti-PD1-Rs experienced a clonal shift back towards the activated profile 2. Importantly, JAKi-Rs began to show a similar clonal shift from profile 9 toward 2, mirroring the transition seen in anti-PD1-Rs, albeit incompletely. This directional shift aligns with clinical improvement in the JAKi-Rs after cycle 4.

A similar dynamic was observed between profiles 1 and 8. NRs showed no consistent transitions across the course of treatment. In contrast, JAKi-Rs exhibited an early shift (cy 1–>2) from profile 1 to profile 8 despite showing no clinical response at this time. Anti-PD1-Rs, on the other hand, did not exhibit such early shifts. Between cycles 4 and 6, both anti-PD1-Rs and JAKi-Rs exhibited a rebound from profile 8 back to profile 1, showing a shared clonal shift in patients responding to therapy.

To interpret these clonal shifts, we examine their transcriptional embeddings. Clones within the above 4 profiles are very plastic, spanning across multiple CD8 T cell subsets (Figure 6E). However, plasticity remarkably declines from profile 1 to 8 and from 2 to 9. For instance, profile 1 clones span CM, EM.GZMK, EFF, and ISG subsets, whereas profile 8 clones concentrate within EFF cells. Similarly, clones in profile 2 span EFF.IFNG and EM.EMRA, and shift toward EMRA.NKG7 in profile 9 (Figure 6F). To systematically quantify the plasticity of a clone, we compute the variance of the cell density distribution in its transcriptomic embedding. As expected, profiles 2 and 9 are enriched with highly plastic clones compared to profile 1 and 8 (Figure 6G), and this trend is consistent across the different response groups (Supplementary Figure 7D). Notably, different response groups show different clone plasticity change during treatment: JAKi-Rs uniquely exhibited a significant decrease in clone plasticity between cycle 1 and cycle 2 that is not seen in the NRs despite having similar early clinical response, while both JAKi-Rs and anti-PD1-Rs showed increased plasticity from cycle 4 to 6 (Figure 6H), consistent with previously identified findings from this study.

Transcriptomic comparisons further revealed that profiles 1 and 2 were characterized by higher expression of activation markers (e.g., *IFNG*, *FOS*, *EGR1*) and lower expression of cytotoxicity markers (e.g., *PRF1*, *GZMB*), compared to profiles 8 and 9 (Supplementary Figure 7E, Supplementary Data 4, Wilcox rank sum test, FDR < 0.05, |logFC| > 0.25). Module score analysis showed that profile 2 had the highest activation score, while profile 9 had the highest cytotoxicity score (Figure 6I, Supplementary Data 5). A similar, though less pronounced, pattern was observed between profiles 1 and 8. Importantly, these functional transitions do not simply reflect differences in cell-type composition: when stratified by cell type, differences in activation and cytotoxicity are observed between the profiles (Figure 6J). Thus, the shifts between profiles 2 and 9 and between profiles 1 and 8 reflect a functional transition from activation to cytotoxicity. These findings are directly compatible with the mechanism of action of anti-PD1 and JAKi respectively. As PD-1 engagement directly blocks TCR-mediated T cell activation via *SHP2* activation, PD-1 blockade increases T cell activation. JAKs are inhibitors of interferon signaling – in particular, type 1 interferon signaling can down-modulate anti-tumor cytotoxic activity^36^; thus, JAKi would be expected to potentially increase cytotoxicity programs.

In summary, projecting temporal information onto the clone embedding revealed clear treatment-associated transitions between profiles 1↔8 and 2↔9, representing temporal changes in plasticity, activation, and cytotoxicity that straddle multiple T cell subsets. The observed plasticity dynamics are consistent with findings from the original study, which reported that JAK1 inhibition modulates T cell fate flexibility and promotes favorable immune states. Importantly, the use of clone profiles in Clonotrace enables detailed characterization of the transcriptional states of responsive clones. Together, our findings suggest that high plasticity and activation— especially during the post-JAKi phase—may underlie favorable clinical outcomes.

## Discussion

Clonotrace introduces a computational framework for integrating molecular and clonal information to resolve cellular dynamics with enhanced precision and interpretability. In this paper, the molecular information is derived from RNA expression and the clonal information is derived from lineage barcodes. Unlike traditional trajectory inference methods that rely solely on transcriptional similarity, Clonotrace leverages clonal structures—whether derived from genetic barcodes or antigen receptor sequences. By modeling each clone as a smoothed density over the cell embedding and quantifying similarity via optimal transport, Clonotrace produces a low-dimensional embedding of the molecular “trace” of each clone to capture the evolution of cells. This approach enables cell fate and pseudotime analysis in challenging scenarios, such as when cell state divisions are subtle or transitory cells are not observed. Clustering within this clonal embedding yields clonotype profiles, which can be projected back onto cells to facilitate downstream tasks such as fate-driving gene detection. By capturing the earliest molecular differences between lineages while adjusting for confounding pseudotime effects, Clonotrace enables rigorous statistical identification of fate driving genes. This strategy is particularly advantageous in the study of immune cells dynamics and therapy-induced plasticity, where fate decisions are jointly influenced by lineage and transcriptional state. Clonotrace further uniquely enables the delineation between unipotent and multipotent fate trajectories.

Across both synthetic and real-world datasets, Clonotrace consistently reveals cellular dynamics that are more informative than what is feasible by transcription alone. We showed this through four examples: In treatment perturbation data, clone-informed pseudotime correctly reconstructed temporal progression when traditional pseudotime methods fail. In lineage barcoded single cell data of hematopoiesis, Clonotrace distinguished bipotent from unipotent clones that were otherwise inseparable in gene expression space. In tumor-infiltrating CD8⁺ T cells, Clonotrace revealed clonal programs linked to cytotoxicity, exhaustion, and memory, identifying fate bifurcations and early markers of differentiation not visible through static clustering. Finally, in longitudinal single cell RNA-seq profiling of peripheral T cells during immunotherapy, Clonotrace identified clonal dynamics associated with clinical outcomes. These findings demonstrate that Clonotrace is a versatile and powerful tool for studying cellular dynamics.

A key innovation of Clonotrace is its clone profile-centric view: rather than relying on predefined cell types or trajectories, it learns recurrent patterns of clonal behavior via optimal transport on the gene expression manifold. This unsupervised framework generalizes to any lineage-resolved single-cell dataset and does not require longitudinal data collection, making it especially suitable for clinical data where temporal resolution is limited.

Despite its strengths, Clonotrace relies on the observation of clone labels and sufficient number of clones with high cell number. Thus, for T cells, Clonotrace can only be applied to systems where there is clonal expansion. When there is not enough clonal expansion, the clonal information adds limited information to a purely transcriptome-based analysis.

Overall, Clonotrace represents a methodological advance for lineage-informed analysis of single-cell data. By bridging clonal identity and transcriptional phenotype, it enables fine-grained dissection of differentiation trajectories, fate potential, and gene regulation. As lineage-resolved single-cell technologies continue to scale, Clonotrace offers a principled approach for interpreting their increasingly rich and multidimensional datasets.

## Methods

### Algorithmic Details of Clonotrace

Clonotrace requires a cell embedding and the cell clone label as input, with 4 main steps as outlined below:

#### 1. Density estimation in the cell embedding for each clone

Clonotrace requires each clone to contain sufficient cell mass to enable accurate estimation of its density distribution in the transcriptomic embedding. We define a threshold of 10 cells per clone to designate expanded clones. However, unassigned cells—those not in expanded clones— provide valuable contextual information, especially if they reside near clonally expanded regions in the transcriptional embedding. To incorporate this contextual information, we infer the probability of an expanded clone having nonzero density at each cell position using label propagation on a cell-cell graph. Let *N* be the total number of cells, *M* be the number of expanded clones, and *Q* ∈ *R*^*N*×*M*^ be the cell-to-clone soft assignment matrix.

We first initialize *Q*. For cell *i* which belong to clone *j*, *j* ∈ {1,2, … , *m*},

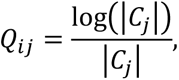

where |*C_i_*| is the number of cells in the clone *j*. For other cells, *Q_ij_* = 0. Such kind of initiation is due to the unbalanced size of clones. Then, *Q* is updated through label propagation: Let W ∈ *R*^*N*×*N*^ be the normalized affinity matrix from the cell-cell graph and let *⍺* ∈ [0,1] be a smoothing parameter. The label propagation updates are as follows:

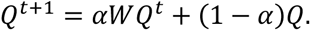

The distribution of cells within expanded clones is not uniform, for example, many cells in early stages (stem cells, T naïve cells) are not in expanded clones. Label propagation in those cell clusters only relies on a few labeled cells, which is likely to introduce false positive label assignment. To filter out the unconfidently assigned labels, we added a bootstrap strategy: in each iteration, the labels from *βN* cells (*β* ∈ (0,1)) would be masked and treated as unlabeled. The label propagation is repeated to get multiple estimations of the cell-to-clone soft assignment. The divergence of the estimation is measured using L1 norm. Cells with divergence higher than the threshold would be excluded. The results of clone label propagation can be viewed in Supplementary Figures 1 and 2.

After label propagation, the cell-to-clone assignment *Q* is thresholded, retaining the top 90% of cumulative probability mass for each cell, setting other probabilities to 0, and then renormalizing the vector.

#### 2. Clone embedding generation

Once we estimate the cell-to-clone assignment probability matrix *Q*, we compare clones based on their smoothed cell density distributions in transcriptomic space using earth mover distance. For each clone, we derive a density distribution across the transcriptomic embedding by normalizing its column in *Q*. Then we build a kNN graph from the cell embedding and define cell–cell distances using shortest path lengths on the graph, which approximates geodesic distances on the manifold approximation of the cell embedding. Then each column in the cell-to-clone probability matrix *Q* can serve as the density distribution for each clone, and this cell-cell geodesic distance can serve as cost matrix to be used to compute the earth mover distance between each pair of clones. The calculation is done by using the ‘transport’ function in the ‘transport’ R package.

The calculation of earth-mover distance based on the above procedure can be computationally prohibitive for large data. As an approximation, for each pair of clones *i* and *j*, we first select the top *x* cells from each clone based on the highest assignment probabilities (if a clone contains fewer than *x* cells, we use all available cells from that clone). For each selected cell in clone *i*, we compute its distance to clone *j* as the average distance to its *k* nearest neighbors among the selected cells in clone *j*, using cell–cell graph distances. The distance from clone *i* to clone *j* is then defined as the average of these cell-to-clone distances. We repeat this process in the reverse direction (from *j* to *i*), and finally define the distance between clones *i* and *j* as the average of the two directional measurements.

This scalable procedure yields a clone-by-clone distance matrix, which can be embedded into a low-dimensional space (e.g., via multidimensional scaling or diffusion maps) to produce the clone embedding. In the analysis of this paper, the clone level pseudotime is inferred in the MDS, and clones are visualized in diffusion maps and UMAP.

#### 3. Clone level pseudotime inference

We adopted the idea of diffusion pseudotime^11^ to infer clone level pseudotime *T* in the MDS projection of the clone-wise distance. Then, the cell-level pseudotime is computed as a weighted average over clone-level pseudotime using the cell-to-clone probability matrix as *QT*.

#### 4. Cell graph reweighting based on clone embedding

To refine the representation of transcriptional relationships based on clonal structure, we reweight the cell–cell graph. In clone embedding (by default the MDS embedding), we build a clone kNN network, the affinity matrix of the clone network is denoted as *V* ∈ *R*^*M*×*M*^. Here we use *D* ∈ *R*^*N*×*N*^ to denote the distance matrix of the cell network, then the reweighted cell distance is generated by *D* ⊘ *QVQ*^*T*^, where ⊘ means element-wise division. Thus, if the clones of two cells are not identical or neighbors in the clone embedding, the distance between the two cells would be inflated.

#### 5. Clone profile generation

Given the clone kNN network we apply Leiden algorithm to cluster clones into *S* profiles. We use *P* ∈ *R*^*M*×*S*^ to denote the clone-to-profile assignment matrix. Then the cell-to-profile probability matrix can be generated by multiplying the cell-to-clone assignment matrix to the clone-to-profile assignment matrix as *QP*.

#### 6. Clone profile enrichment test within a transcriptionally defined cell subset

To test whether certain clonotype profiles are enriched in specific cell types, we first constructed a contingency table recording the mass of cells assigned to each clonotype profile within each cell type. To assess enrichment, we performed a permutation test by randomly shuffling the cell type labels and recalculating the profile-by-cell-type distributions for each permutation. Finally, we compute empirical p-values for observed enrichments.

#### 7. Fate driving genes detection

To identify genes associated with early fate bias, within the target transcriptomic cluster containing multiple profiles, we model expression for gene *g* in cell *i* in profile *s*, *s* = [1, … , *S*] as a function of pseudotime *t*_*i*_: *g*(*t*_*i*_)_*s*_ = *B*(*t*_*i*_)*β*_*s*_, *B*(*t*) is the spline function with the degree of freedom denoted by *df*. *β*_*s*_ are specific to each profile. Given the cell-to-profile soft assignment matrix *P* ∈ *R*^*N*×*S*^, the expression of gene *g* in cell *i* in the target cluster can be modeled as: 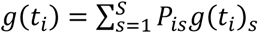. To test if the targeted profile *a* has a different *β*_*a*_ compared to other profiles, we aggregate the probabilities for all other profiles as control profile *b*, so the hypothesis testing is framed as:

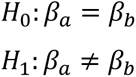

Then we do F-test to determine if any profile has a different gene expression pattern along the pseudotime axis. In practice, within a cell cluster there is usually a short time frame, within which we assume the gene dynamics won’t be complex, thus we set *df* = 1 for the spline fit, which is a linear regression.

### Data Simulation

Data was simulated for Figure 2 based on the following workflow.

#### 1. Cell assignment to profiles

The simulations are generated from real single cell transcriptomics data. Cell trajectories are first generated by spline fitting across specified clusters. Trajectories are then manually assigned to simulated profiles. Then cells are softly assigned to profiles given the minimum distance to the trajectories. Taking the first simulation as an example, in the bipotent profile there are two trajectories (named with *x* and *y*) and the unipotent profile has one trajectory (named with *z*). Then we calculate the distance between each cell and each trajectory. Then the probability for cell *i* to be assigned to the bipotent profile is

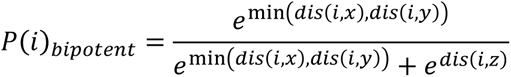

And for the unipotent profile:

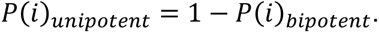

#### 2. Cell time simulation

Each trajectory is simulated with a time range. Time range always starts from 0 but can have different end points. For example, the time range for trajectory *j* is [0, 0.8], which means the cell at the end of this trajectory would be assigned with time as 0.8. Then we generate the time for each cell *i* using the weighted average of time projection on each trajectory:

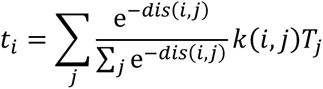

*j* denotes the id of trajectories. *dis*(*i*, *j*) means the distance between cell *i* and trajectory *j*. *k*(*i*, *j*) denotes the relative position that cell *i* projected on trajectory *j*. And *T*_i_ means the upper bound of the time range.

#### 3. Clone generation

Given the cell to profile probability, each cell is randomly assigned with a profile label. Then within each profile, we generate clones by iterative sampling without replacement. The cell sampling for each clone is guided by the simulated time generated in step 2. Each time a cell would be chosen as the center of a clone. Using *t* to denote the simulated time for the center cell, the clone size is specified by one-time sampling from exponential distribution with 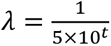. Then cells are sampled to this clone with the probability as *normal*(*t*, 0.1*t* + 0.05). Thus, this simulation process assumes each clone to expand and become more diverse along time. Once cells are sampled to a clone, those cells would be removed from the candidate pool to avoid repetitive sampling. The iteration stops when there are not enough cells to satisfy the size requirement for a clone.

### Baseline for benchmark in simulation

Within the target cluster, cells are classified into different profiles given if their clones extended to the target terminal fates. The comparison of the gene expression between different profiles is done by Wilcoxon’s rank sum test. To account for the different distributions along the time, we identified top *k* mutual nearest neighbors between profiles, and cells without mutual nearest neighbors would be filtered out in the downstream test.

### Module score calculation

The module scores are calculated using the *AddModuleScore* function from the Seurat package (v5.0.1) based on provided genes lists for different functions.

### Cdr3b distance calculation

The cdr3b distance for expanded clones is calculated using the tcrdist python package (v 0.2.2) with the default parameters. The reference database is ‘alphabeta_gammadelta_db.tsv’.

### Clone plasticity calculation

To quantify the transcriptional plasticity of each clone, we compute the total variance of cells associated with that clone, using the soft assignment probabilities as weights. Let X ∈ *R*^*N*×*D*^denote the matrix of cell embedding, where *N* is the number of cells and *D* is the feature dimension. *Q* ∈ *R*^*N*×*M*^is the cell-to-clone soft assignment matrix, where *P*_*ik*_ represents the probability that cell *i* belongs to profile *k*. For each clone *k* ∈ [1, … , *M*], the assignment weights are normalized as

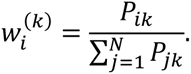

We then compute the weighted mean ***μ***^(*k*)^ ∈ *R*^*D*^ with 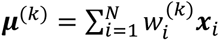, in which *x*^*i*^ ∈ *R*^*D*^ is the coordinate vector of cell *i* in the cell embedding. Next, we calculate the weighted covariance matrix 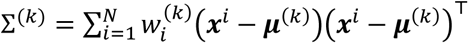. And we define clone plasticity for profile *k* as the trace of the covariance matrix: 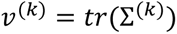.

## Acknowledgements

This research was supported by National Institutes of Health Grant 5R01HG006137 (NRZ), 5R01GM149671 (NRZ), DMS/NIGMS grant DMS-2245575 (NRZ), 1R56AG081351 (NRZ), F32NS108580 (CJ),Neurosurgery Research Education Foundation (CJ), Funding through the Bloomberg∼Kimmel Institute for Cancer Immunotherapy (DP), and through Mark Foundation Center for Immunotherapy, Immune Signaling, and Radiation (NRZ).

## Contributions

Conceptualization: Y.F., N.R.Z., C.J.

Initial data analysis and preprocessing: Y.F., C.J.

Algorithm for Clonotrace: Y.F.

Algorithm development and implementation: Y.F.

Simulation design and data analysis: Y.F.

Cancer treatment sample generation: D.S

Data preprocessing for cancer treatment sample: X.Y.C, K.Z.L

Data analysis: Y.F.

Benchmarking: Y.F.

Manuscript writing: Y.F., N.R.Z. with feedback from D.M., M.W., C.J., D.P., X.Y.C, S.M.S

Supervision: N.R.Z.

## Data availability

The longitudinal single cell RNA-seq for hematopoiesis is publicly available from the Gene Expression Omnibus with the accession GSE140802 (https://www.ncbi.nlm.nih.gov/geo/query/acc.cgi?acc=GSE140802). The single cell RNA-seq for the treated cancer cell lines is waiting to be published in Gene Expression Omnibus recently. The single cell RNA-seq for PBMC T cells from NSCLC patients is publicly available from the Gene Expression Omnibus with the accession GSE266219 (https://www.ncbi.nlm.nih.gov/geo/query/acc.cgi?acc=GSE266219). The single cell RNA-seq for the treated cancer cell lines and the single cell RNA- and TCR-seq for tumor infiltrating T cells in glioblastoma will be made available in Gene Expression Omnibus upon publication.

## Code Availability Statement

The scripts for the analysis are wrapped into the R package Clonotrace: https://github.com/yuntianf/Clonotrace.

## Notes

https://github.com/yuntianf/Clonotrace

### Competing Interest Statement

The authors have declared no competing interest.

