## Supplementary Figures for "Deciphering Cell Fate and Clonal Dynamics via Integrative Single-Cell Lineage Modeling"

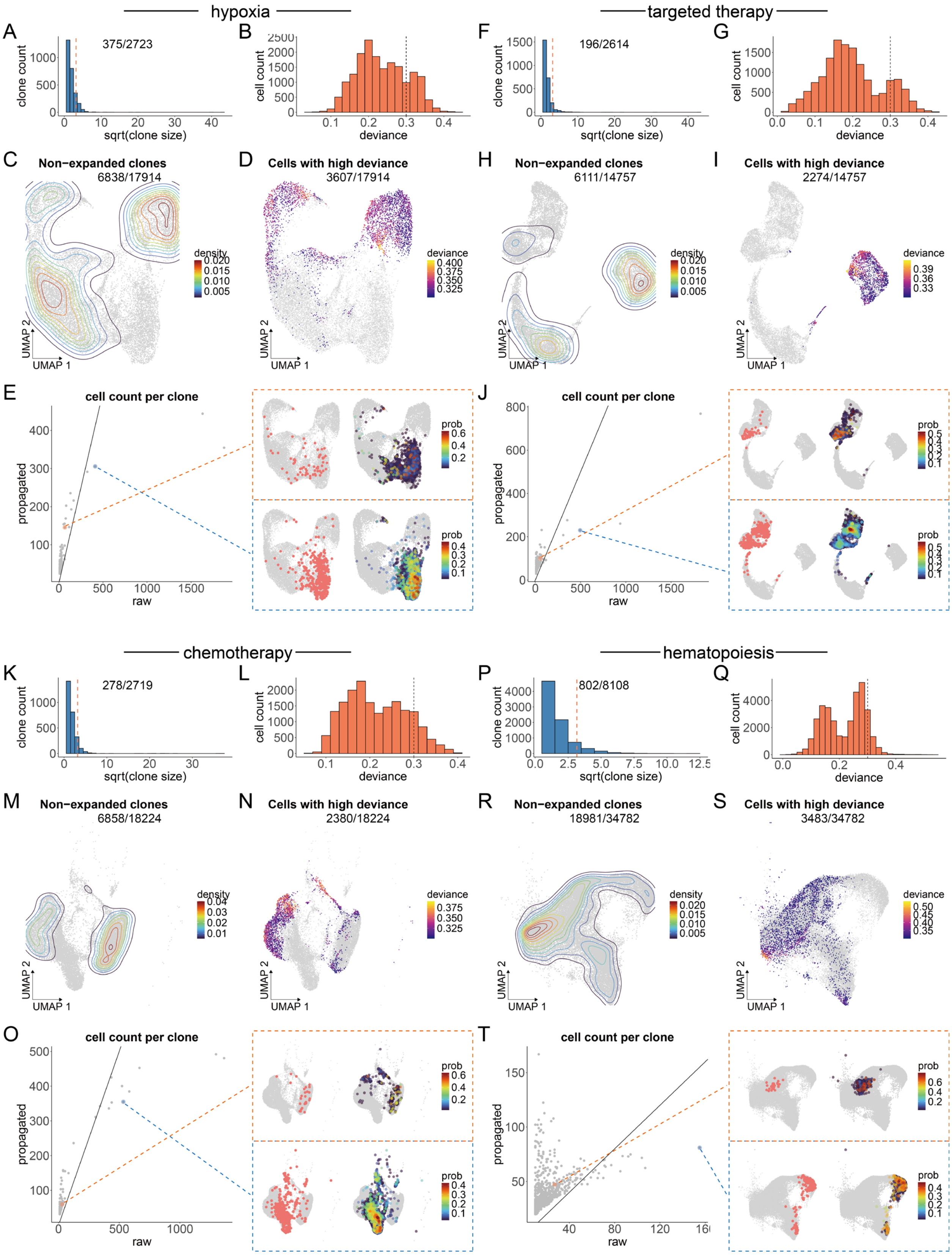


**Supplementary Figure 1 : Clone label spreading in the treated cancer cell lines and**

**hematopoiesis. A** Histogram for the clone size distribution for cancer cells treated with hypoxia. Clone size = 10 is highlighted with a red dashed line. The number of expanded clones (size >= 10) out of the total number of clones is marked at the top. **B** Histogram of the deviance for cells during the bootstrap of clone label spreading. The threshold (deviance < 0.3) is highlighted with a black dashed line. **C** Cell density distribution for cells within non-expanded clones over the cell transcriptomic UMAP (clone size < 10). The number of cells within non-expanded clones out of all cells is marked at the top. **D** The distribution for cells with high deviance (> 0.3) during clone label spreading over the cell transcriptomic UMAP. Cells are colored by deviance. **D** Distribution for cells with high deviance (> 0.3) during clone label spreading over the cell transcriptomic UMAP. Cells are colored by deviance. **E** Comparison of the cell mass before and after clone label spreading for expanded clones. In the left scatter plot, each point is a clone. The line denotes the equivalence. Two example clones are highlighted in coral and blue. The cell density distribution of the two example clones before and after label spreading are shown in the right UMAPs. **F-J** are formatted the same as **A-E** to show the clone label spreading for cancer cell treated with targeted therapy. **K-O** are for cancer cells treated with chemotherapy. **P-T** are for cells in hematopoiesis.

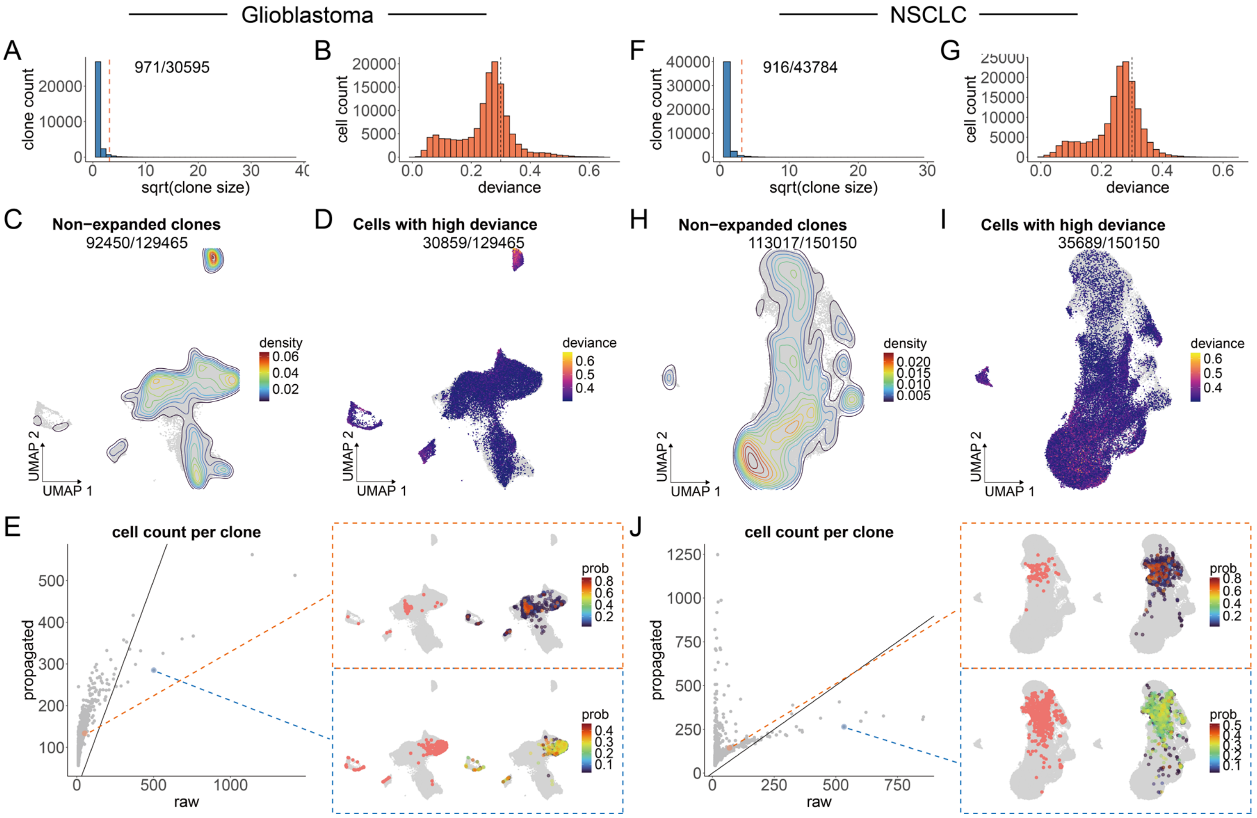


**Supplementary Figure 2: Clone label spreading in the tumor infiltrating T cells in glioblastoma and PBMC cells in patients with NSCLC. A-E** are formatted the same as **Supplementary Figure 1 A-E** to show the clone label spreading for tumor infiltrating T cells in glioblastoma. **F-J** are for cells in PBMC in patients with NSCLC.

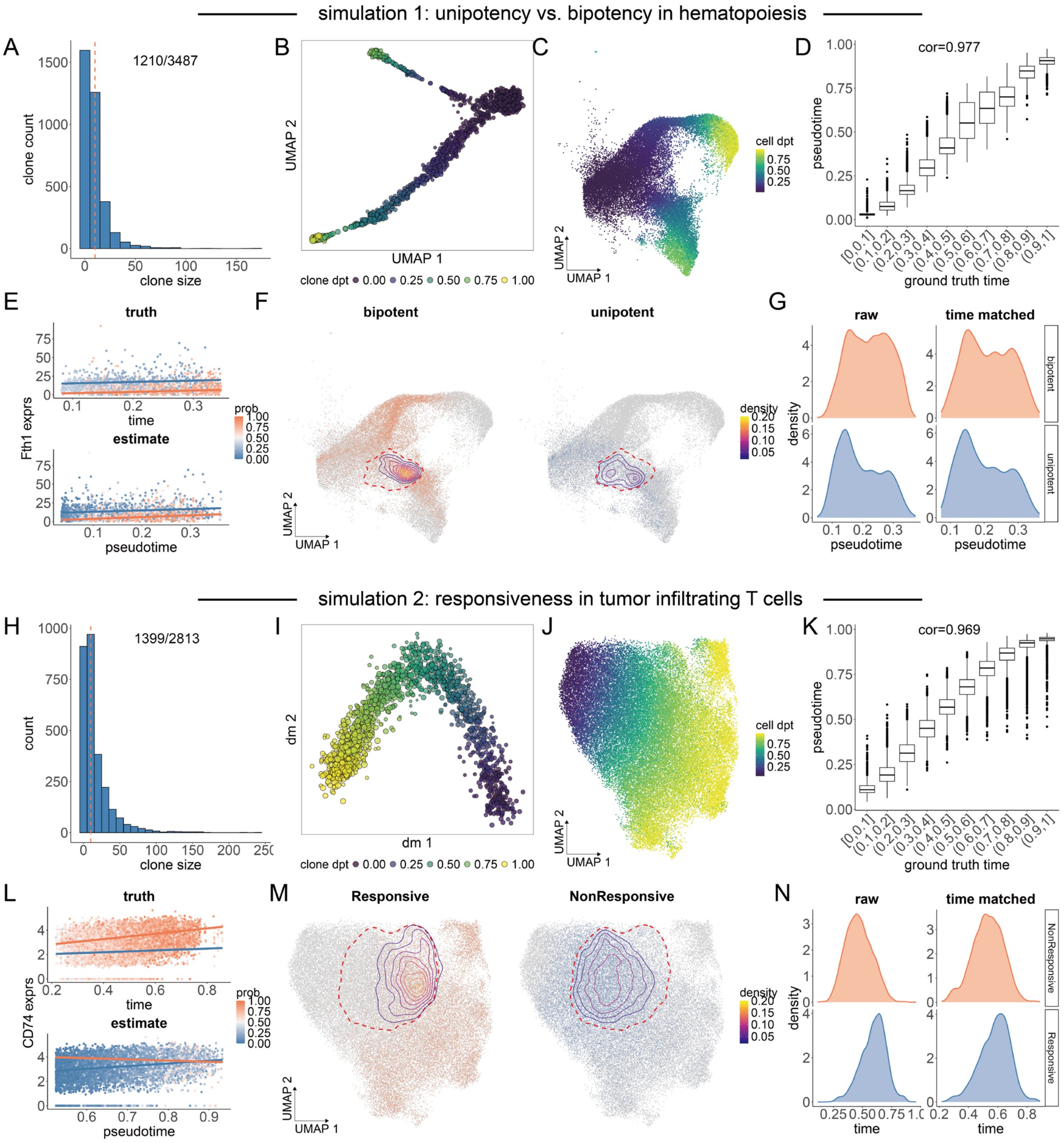


**Supplementary Figure 3: Pseudotime inference and scheme for fate driving gene detection in the simulations. A** The clone size distribution in the simulated data based on the hematopoiesis. Clone size = 10 is highlighted with a red dashed line. The number of expanded clones (size >= 10) out of the total number of clones is marked at the top. **B** Clone level diffusion pseudotime. Each point is a clone colored by the inferred pseudotime. **C** Smoothed clone level diffusion pseudotime in the cell embedding. Each point is a cell colored by the smoothed pseudotime. **D** Comparison between the ground-truth time in the simulation and the inferred pseudotime. The Pearson correlation between the ground truth and the inference is marked at the top. **E** Example of fate-driving gene detection using linear regression in the ground-truth (top) and Clonotrace-inferred (bottom) settings. Each point is a cell colored by its profile probability; regression lines for bipotent (coral) and unipotent (blue) cells are overlaid. **F** The earliest separation between the bipotent and unipotent lineage (cluster 4) is highlighted in red dashed circle. Left: Cells whose clones span clusters 1, 6, or 7 (bipotent progenitors) are colored coral. Right: Cells whose clones are restricted to clusters 3 or 5 (unipotent progenitors) are colored blue. Contours indicate cell density in the cluster 4. **G** Distribution of bipotent and unipotent progenitors along simulated time within cluster 4, before (left) and after (right) time matching. **H-N** are formatted the same as **A-G** for the simulation based on tumor infiltrating T cells in glioblastoma.

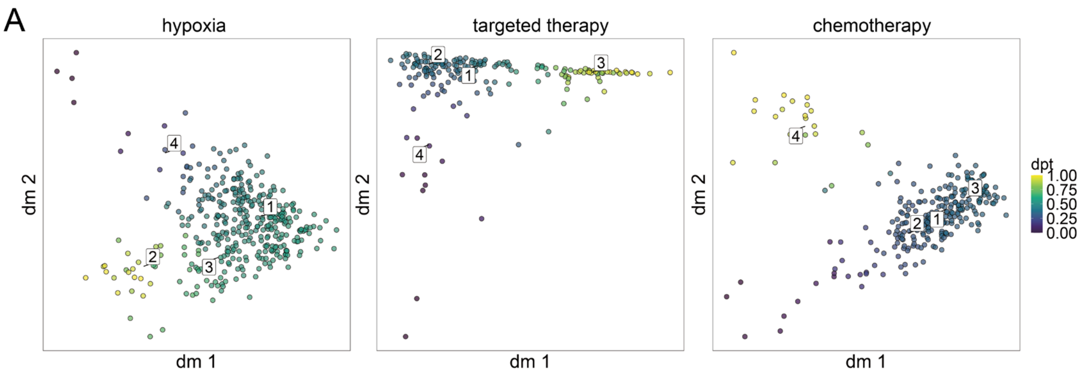


**Supplementary Figure 4: Clone level pseudotime for cancers treated with different therapies. A** The diffusion map for the clone embedding of cancers treated with hypoxia (left), targeted therapy (middle) and chemotherapy (right). Each dot is a clone colored by the clone level pseudotime.

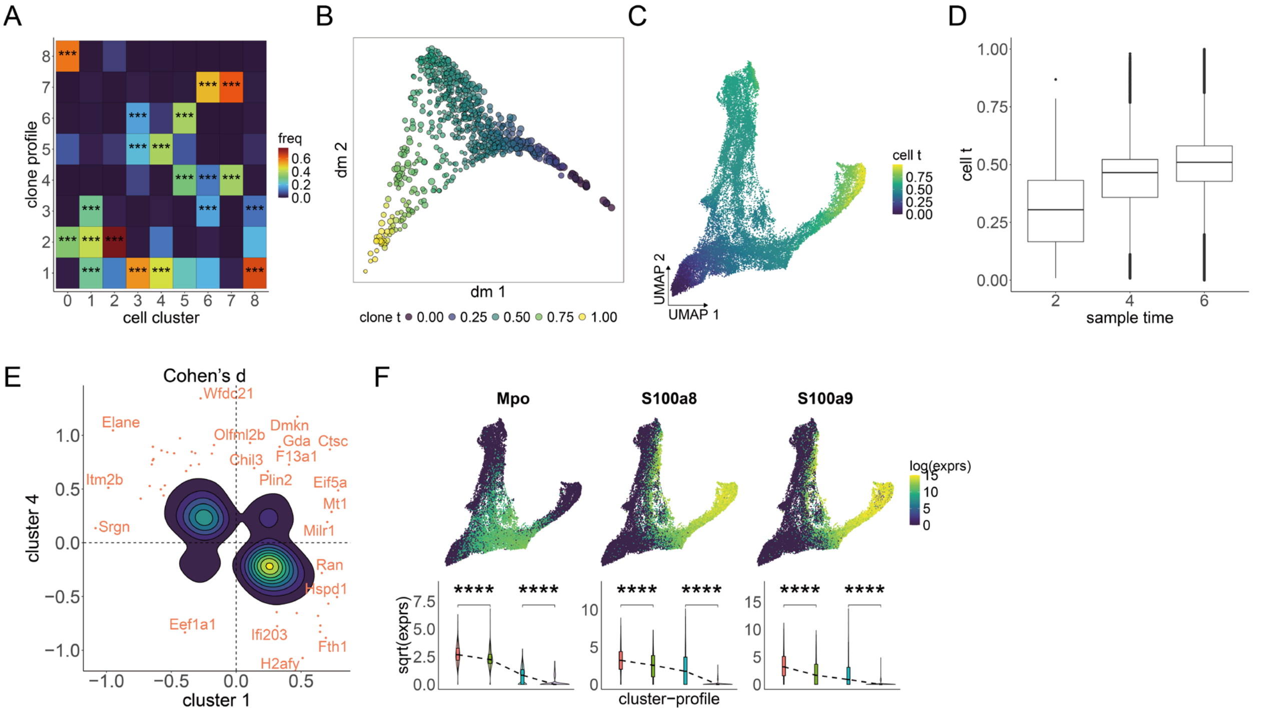


**Supplementary Figure 5: Supplements for the hematopoietic data analysis. A** Clone profile enrichment in each cell cluster. The color of each tile indicates the frequency of cells for the given profile in the given cell type (normalized within each cell type). Significant enrichments are highlighted with *. **B** Clone level diffusion pseudotime. Each point is a clone colored by the inferred pseudotime. **C** Smoothed clone level diffusion pseudotime in the cell UMAP. Each point is a cell colored by the smoothed pseudotime. **D** Comparison between the sample time and the inferred pseudotime. **E** Distribution of Cohen's d for significant genes for profile 1 in both cluster 1 and 4. The contour shows that most significant genes show inverse direction of Cohen's d, indicating an intermediate expression of those genes in profile 1. Genes with high effect size are highlighted in coral points with gene names. **F** Expression distribution for *Mpo* and *S100a8/a9*. Top, the log-wise expression for example genes over the cell UMAP. Bottom, the comparison of gene expression (transformed by square-root) between the three profiles in cluster 1 and 4.

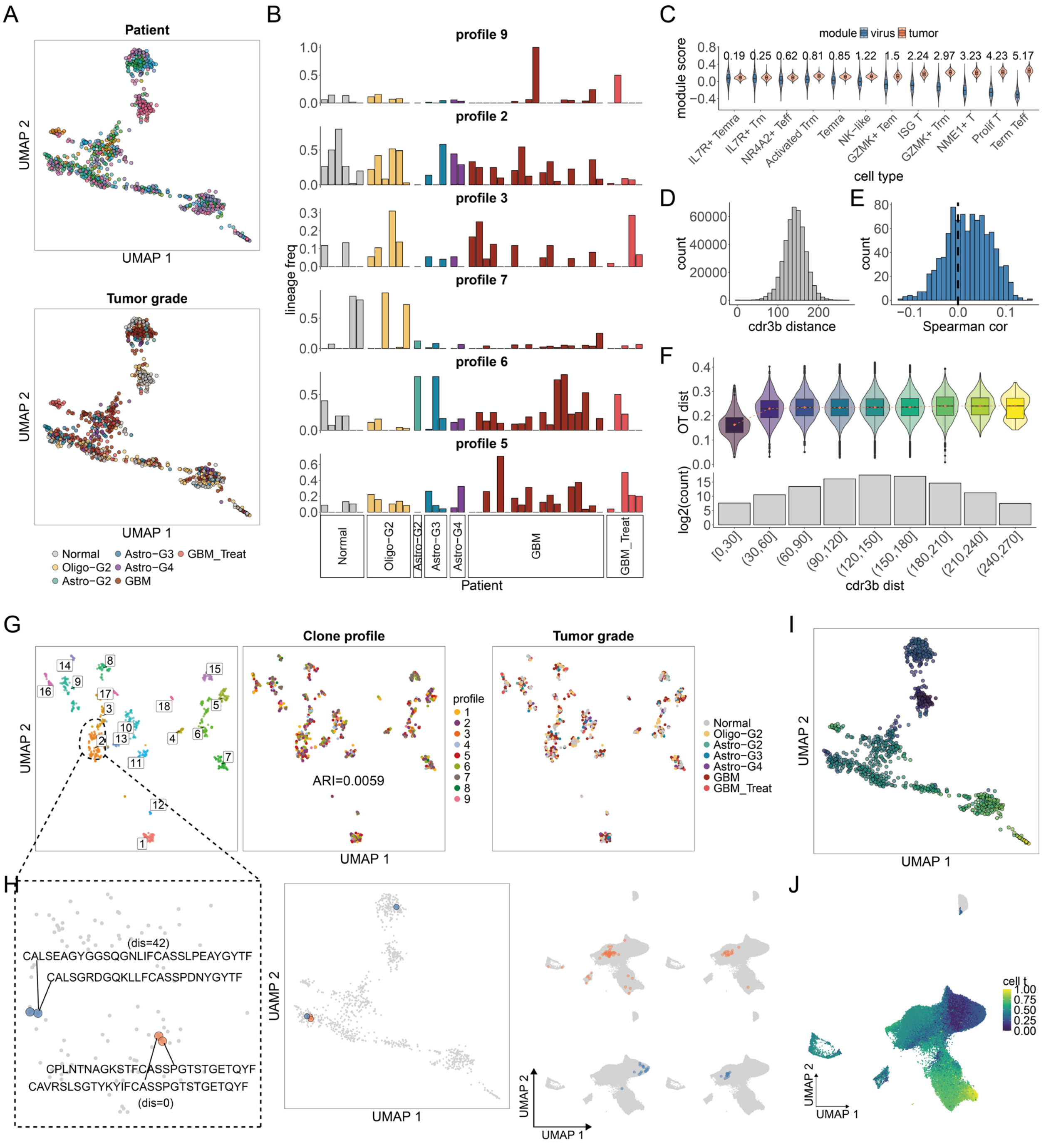

**Supplementary Figure 6: Supplements for the analysis of tumor infiltrating T cells in glioblastoma. A** Distribution of patients and tumor grades over the clone UMAP. Each point is a clone colored by patient identity (top) or the tumor grade (bottom). Each profile covers multiple patients. **B** Clone distribution in different patients across tumor grades in the left profiles (2, 3, 5, 6, 7 and 9). Patients are ordered and colored by tumor grade. The height of the bar denotes the frequency of clones in the given profile for the patient. **C** Module score for tumor reactiveness (coral) and virus reactiveness (blue) in each cell type. **D** Distribution of the cdr3b distance between expanded clones (clone size >= 10). **E** Distribution of Spearman correlation between cdr3b distance and clone transcriptomic distance. The position where Spearman correlation is 0 is marked with a black dashed line. **F** Top: distribution of clone transcriptomic distance across the bins of cdr3b distance. Violins are colored by the cdr3b distance. Bottom: The number of distance pairs in each cdr3b distance bin. **G** UMAP projected from the cdr3b pairwise distance. Each point is a clone colored by cdr3b cluster (left), clone profile (middle) and tumor grade (right). The adjusted rand index between the cdr3b cluster and clone profile is marked at the center. **H** Clone examples to show the low correlation between cdr3b sequence and transcriptomic distribution. Left: zoom-in of the cluster 2 from the UMAP based on cdr3b distance. The two clone example pairs are highlighted with coral and blue. The coral pair has very low cdr3b distance while the blue pair has a slightly higher distance. The corresponding cdr3b sequences and cdr3b distance are marked.  Middle: The location of the clone example pairs in the clone transcriptomic embedding. Though two blue clones have similar cdr3b sequences, they have very different cell distribution in the transcriptomic embedding. Right: the cell distribution of the two clone example pairs. **I** Clone level diffusion pseudotime. Each point is a clone colored by the inferred pseudotime. **J** Smoothed clone level diffusion pseudotime in the cell UMAP. Each point is a cell colored by the smoothed pseudotime.

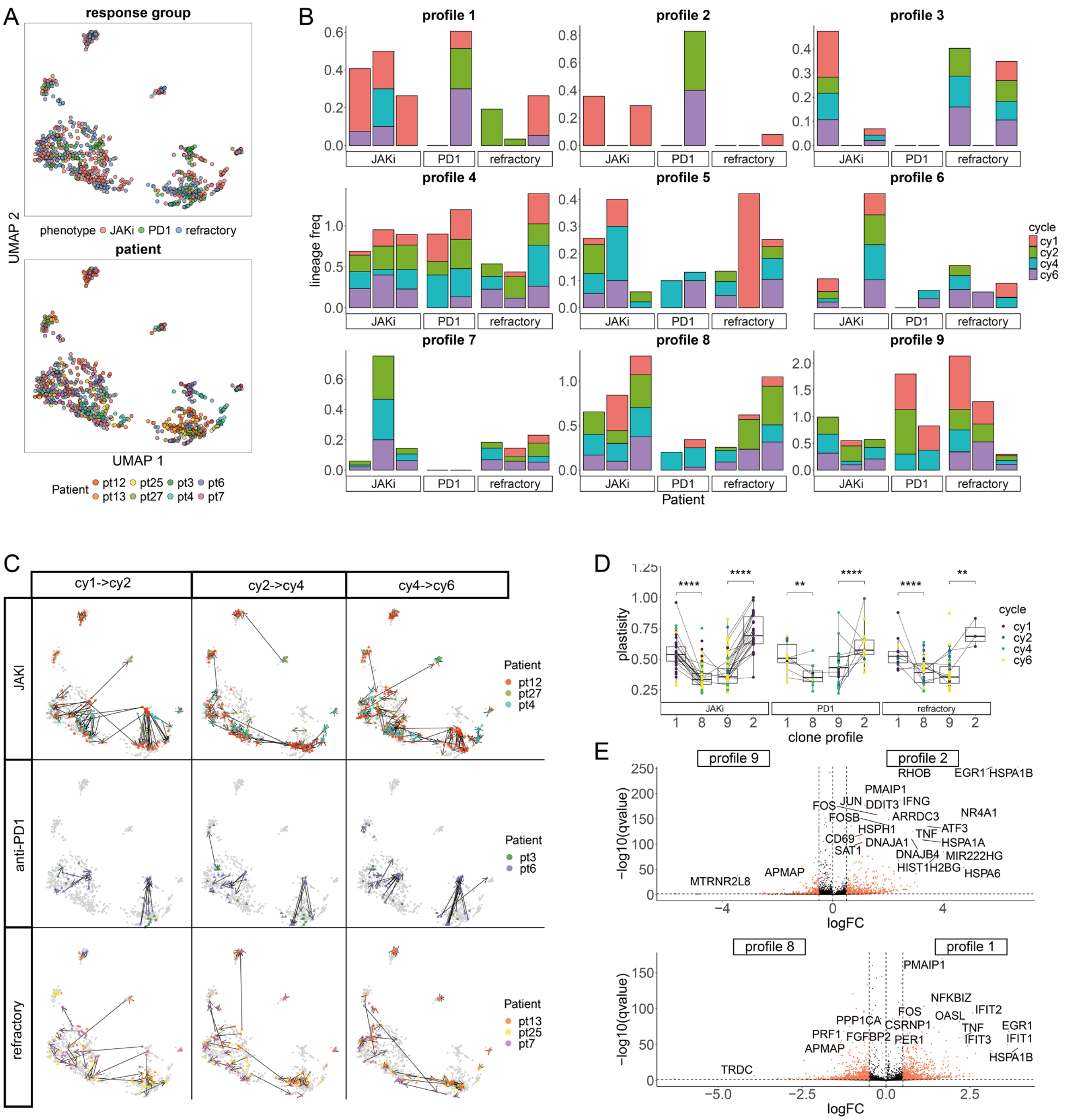


**Supplementary Figure 7 : Supplements for the analysis of PBMC T cells from patients with NSCLC. A** Distribution of patients and tumor grades over the clone UMAP. Each point is a clone colored by response phenotype (top) and the patient identity (bottom). Each profile covers multiple patients. **B** Clone distribution in different patients across tumor grades in each profile. Patients are ordered by response phenotype in the x axis. The height of the bar denotes the frequency of clones in the given profile for the patient. Time points are represented by different colors. **C** Clone transition during the treatment across all profiles. Each row is a response group, and each column is a treatment cycle. Each point is a clone colored by the patient source.

Clones with the same TCR at different cycles are connected by an arrow. **D** The shifts of clone plasticity across treatment cycles in each patient group. Each point is a clone colored by treatment cycle. Clones with the same TCR at different time points are linked by a line. Significant changes in plasticity between profiles are highlighted with *. **E** Volcano plots to show the differentially expressed genes between profile 9 and 2 (top) and profile 1 and 8 (bottom). Each point is a genes, significant genes with high effect size (fdr < 0.05, Cohen's d > 0.5) are highlighted in coral with gene names. Fdr=0.05, Cohen's d = 0.5 or -0.5 are marked with black dashed lines.
